# Profiling acetogenic community dynamics in anaerobic digesters - comparative analyses using next-generation sequencing and T-RFLP

**DOI:** 10.1101/2021.01.26.427894

**Authors:** Abhijeet Singh, Bettina Müller, Anna Schnürer

**Author notes:** For correspondence.; Address: Department of Molecular Sciences, Box 7025, 75007 Uppsala, Sweden.

## Abstract

Acetogens play a key role in anaerobic degradation of organic material and in maintaining biogas process efficiency. Profiling this community and its temporal changes can help evaluate process stability and function, especially under disturbance/stress conditions, and avoid complete process failure. The formyltetrahydrofolate synthetase (FTHFS) gene can be used as a marker for acetogenic community profiling in diverse environments. In this study, we developed a new high-throughput FTHFS gene sequencing method for acetogenic community profiling and compared it with conventional T-RFLP of the FTHFS gene, 16S rRNA gene-based profiling of the whole bacterial community, and indirect analysis via 16S rRNA profiling of the FTHFS gene-harbouring community. Analyses and method comparisons were made using samples from two laboratory-scale biogas processes, one operated under stable control and one exposed to controlled overloading disturbance. Comparative analysis revealed satisfactory detection of the bacterial community and its changes for all methods, but with some differences in resolution and taxonomic identification. FTHFS gene sequencing was found to be the most suitable and reliable method to study acetogenic communities. These results pave the way for community profiling in various biogas processes and in other environments where the dynamics of acetogenic bacteria have not been well studied.

**GRAPHICAL ABSTRACT:** 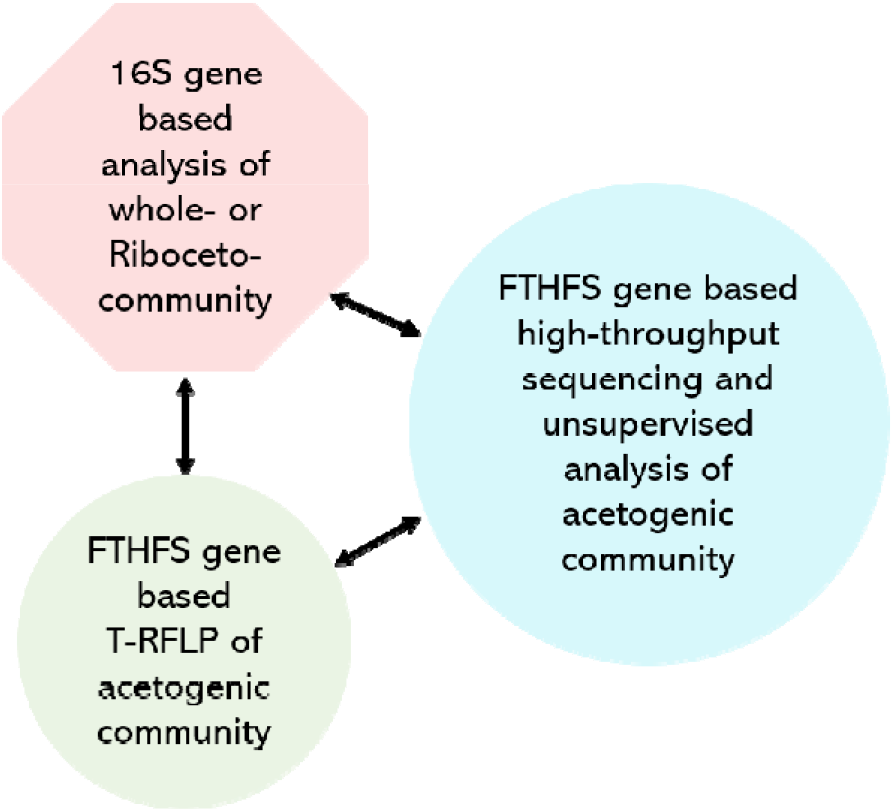

**ONE SENTENCE SUMMARY:** Our high-throughput FTHFS gene AmpSeq method for barcoded samples and unsupervised analysis with AcetoScan accurately reveals temporal dynamics of acetogenic community structure in anaerobic digesters.

## INTRODUCTION

Anaerobic digestion (AD) is a microbiological process through which almost any biodegradable material can be transformed into renewable biofertiliser and biogas, which is mainly a mixture of methane (60-70%) and carbon dioxide (30-40%) (Petersson and Wellinger 2009; SGC 2012; Ma, Yin and Liu 2017; Ruan *et al.* 2019). Anaerobic digestion technology currently serves the purpose of carbon recycling of various waste streams via biogas and organic fertiliser, but also has immense potential in alleviating climate change and hypertrophication (Scarlat, Dallemand and Fahl 2018; Winquist *et al.* 2019). The amount and composition of the biogas produced, and the efficiency and stability of the process, are influenced by various parameters such as feedstock composition, digester technology, operating parameters and the composition and activity of the microbiological community engaged in the process (Pöschl, Ward and Owende 2010; Angelidaki *et al.* 2011; Herrmann *et al.* 2012; Wellinger, Murphy and Baxter 2013; Lebuhn *et al.* 2015; Horváth *et al.* 2016; Schnürer and Jarvis 2017).

The AD process comprises four major complex and interrelated microbiological steps (hydrolysis, acidogenesis, anaerobic oxidation, methanogenesis) carried out by a complex microbial community composed of archaea, obligate and facultative anaerobic bacteria and anaerobic fungi (Zhou *et al.* 2002; Hattori 2008; Thauer *et al.* 2008; Angelidaki *et al.* 2011; Dollhofer *et al.* 2015; Schnürer 2016; Vinzelj *et al.* 2020). Acetogenic bacteria play a critical role in the AD process by performing both reductive acetogenesis and syntrophic oxidation of organic acids, and thus act as a vital link between the hydrolysing and fermenting microbial community and the methanogenic archaea (Ryan, Forbes and Colleran 2008; Ryan *et al.* 2010; Hori *et al.* 2011; Ivarsson *et al.* 2016; Drake *et al.* 2017; Williams, Joblin and Fonty 2020). The overall community is influenced by many parameters, *e.g.* organic substrate, hydraulic retention time (HRT), organic loading rate (OLR) and temperature (Sun *et al.* 2014; Moestedt *et al.* 2016). For an efficient biogas process, different microbiological steps must be balanced and synchronised, otherwise the process can experience disturbance and accumulation of degradation intermediates such as volatile fatty acids (VFA) (Schnürer 2016; Schnürer and Jarvis 2017). One strong regulating parameter for the microbial community is the ammonium/ammonia level, set by the substrate and operating conditions (De Vrieze *et al.* 2015; Robles *et al.* 2018). High levels of free ammonia often result in significant inhibition of methanogenesis and sometimes also of hydrolysis and fermentation (Siegert and Banks 2005; Wang *et al.* 2009; Franke-Whittle *et al.* 2014; Schnürer 2016; Westerholm, Moestedt and Schnürer 2016; Schnürer and Jarvis 2017; Czatzkowska *et al.* 2020). As a consequence, ammonia inhibition also results in accumulation of VFA, particularly propionate, which can further enhance inhibition, cause process instability and reduced methane production (Schnürer and Nordberg 2008; Rajagopal, Massé and Singh 2013; Frank *et al.* 2016; Moestedt *et al.* 2016; Schnürer 2016). Under such stress conditions, acetogenesis/syntrophic acetate oxidation involving specialist methanogens (less sensitive towards ammonia) for methane production is the major pathway which drives the process forward (towards methanogenesis) (Hori *et al.* 2011; Westerholm *et al.* 2011; Schnürer 2016; Westerholm, Moestedt and Schnürer 2016). Conclusively, the acetogenic community can be a good marker when monitoring the health and stability of the biogas process (Hattori 2008; Müller *et al.* 2016; Singh *et al.* 2020).

In recent years, various molecular biological techniques have been applied to investigate and understand the composition, structure and dynamics of the AD microbiome and its implications for the biogas process (Cabezas *et al.* 2015; Lebuhn *et al.* 2015; Schnürer 2016). Advanced and accurate meta-omics technologies (metagenomics, metaproteomics, metatranscriptomics, metabolomics) can help resolve the phylogeny, interactions and functions of microbial species (Vanwonterghem *et al.* 2016). However, these methodologies are generally used to obtain snapshots of the microbial community (Prosser 2015) and are less practical (too expensive, laborious and resource-intensive) for tracking the temporal dynamics over extended periods (Greninger 2018; Martin *et al.* 2018). Thus, less expensive and relatively easier molecular marker-based analysis techniques are generally used to track the temporal dynamics of the whole microbial community involved in the biogas process or a selected fraction of that community. These techniques include fluorescence *in-situ* hybridisation (FISH), single-strand conformation polymorphism (SSCP), denaturing gradient gel electrophoresis (DGGE), quantitative real-time polymerase chain reaction (qRT-PCR), terminal restriction fragment length polymorphism (T-RFLP) and amplicon sequencing (AmpSeq) of the 16S rRNA gene (Čater, Fanedl and Logar 2013; Robles *et al.* 2018). However, few studies have specifically focused on acetogenic community dynamics or on developing a high-throughput method for reliable acetogenic community profiling (Singh *et al.* 2020). Since acetogenesis is a physiological process, and not a phylogenetic characteristic, development of acetogen-specific 16S rRNA gene primers is practically impossible (Ljungdahl 1986; Drake 1994a; Lovell 1994; Lovell and Leaphart 2005; Drake, Gößner and Daniel 2008; Singh *et al.* 2019, 2020). Therefore, established molecular analysis methods/pipelines based on the 16S rRNA gene cannot be used for analysis of the acetogenic community. The Wood-Ljungdahl pathway (WLP) is a characteristic of acetogens (Drake 1994b; Peretó *et al.* 1999; Lever 2012; Poehlein *et al.* 2012) and the marker gene formyltetrahydrofolate synthetase (FTHFS) has been successfully used to decode the potential acetogenic community (Hori *et al.* 2011; De Vrieze and Verstraete 2016; Müller *et al.* 2016). While FTHFS is also present in the genome of non-acetogenic bacteria, sulphate-reducing bacteria and methanogens, still it has been successfully used for over two decades as the marker of choice in acetogenic community analysis (Ljungdahl 1986; Lovell, Przybyla and Ljungdahl 1990; Lovell and Leaphart 2005; Ohashi *et al.* 2007; Drake, Gößner and Daniel 2008; Gagen *et al.* 2010; Moestedt *et al.* 2016; Schuchmann and Müller 2016; Singh *et al.* 2019, 2020). Recently, we published a database (AcetoBase) (Singh *et al.* 2019) and an analysis pipeline (AcetoScan) (Singh *et al.* 2020), and successfully demonstrated proof-of-concept for targeting the potential acetogenic community in biogas reactor samples and unsupervised analysis of FTHFS AmpSeq data. Our previous studies showed that AcetoBase and AcetoScan can be used for reliable monitoring of the acetogenic community in multiplexed samples in biogas reactors.

The aims of the present study were to 1) further evaluate AcetoBase and AcetoScan for profiling and monitoring the temporal dynamics of acetogenic communities in biogas reactors and 2) compare this new high-throughput AmpSeq method targeting the acetogenic community with conventional methods such as T-RFLP and 16S rRNA gene sequencing. Specifically, three different methods (16S rRNA gene AmpSeq, T-RFLP and AmpSeq of FTHFS gene) were evaluated for their ability to monitor the dynamics of the potential acetogenic community in two laboratory-scale biogas processes. In addition, the FTHFS gene-harbouring community was analysed indirectly by profiling the corresponding16S rRNA genes using the 16S rRNA database RibocetoBase (this study), which was deduced from the Silva rRNA database (Quast *et al.* 2013). The selected biogas reactors were operated with food waste under high-ammonia conditions. Method comparisons were performed using samples from a stable control reactor and a reactor exposed to controlled overloading resulting in instability, followed by a recovery phase. Samples from these biogas digesters were used because the dynamics of both the acetogenic and syntrophic acetate-oxidising bacterial community had been identified in our previous studies, enabling comparative analysis (Westerholm *et al.* 2015; Müller *et al.* 2016; Singh *et al.* 2020). We evaluated the overall potency and utility of the different methods by visualisation of potential acetogenic community dynamics in a biogas environment.

## MATERIALS AND METHODS

### Sample collection and processing

Samples were collected in a time-series manner (Supp. data 1) from two parallel mesophilic (37 °C) continuously stirred-tank biogas reactors (active volume 5 L), denoted GR1 (experimental) and GR2 (control), in the Anaerobic Microbiology and Biotechnology Laboratory, Swedish University of Agricultural Sciences, Uppsala, Sweden. Both reactors were operated with mixed food waste at an OLR of ~2.5 g volatile solids (VS) L^-1^ day^-1^, ammonium-nitrogen (NH_4_^+^-N) ~5.4 g/L (free ammonia (NH_3_) 0.6-0.9 g/L) and HRT 30 days, while other operating parameters were as described previously (for reactor D^TE^37 (Westerholm *et al.* 2015). Before the start of the experiment, both reactors had a stable carbon dioxide (32-35%) and methane content (58-60%). To disrupt the stable microbial community, a controlled overloading disturbance was induced in GR1 by increasing the OLR to ~4.09 g VS L^-1^ day^-1^, while control reactor GR2 continued at OLR ~2.5 g VS L^-1^ day^-1^. The increase in organic load (Δ ~1.59 g VS L^-1^ day^-1^) marked the start of the experiment (day 0). The first sample was taken one day after the start of the experiment and the reactors were operated for 350 days with subsequent sampling. Monitoring was based on gas composition and total VFA in the reactors, using GR2 as reference. The OLR for GR1 was returned to ~2.5 g VS L^-1^ day^-1^ when the carbon dioxide content in biogas was observed to be ~50 % (at ~125 days). Extraction of genomic DNA from the samples was performed in triplicate using the FastDNA™ Spin kit for soil (MP Biomedicals), with an additional wash step with 5.5 M guanidinium thiocyanate (Sigma-Aldrich 2020) for humic acid removal (Singh 2020a). Samples and extracted DNA were stored at −20 °C until further use.

### Experimental T-RFLP library preparation and data analysis

Sixteen samples from different time points (Fig. 1; Supp. data 1) were used for T-RFLP profiling and partial FTHFS gene amplicons were generated by the primer pairs and PCR protocol developed in our previous study (Müller, Sun and Schnürer 2013), with the modifications of FAM labelled FTHFS_fwd (5’-FAM-CCIACICCISYIGGNGARGGNAA-3’) and non-labelled FTHFS_rev (5’-ATITTIGCIAAIGGNCCNCCNTG-3’). The FTHFS amplicons were purified by E-Gel^®^ iBase™ Power System (Invitrogen 2012) and E-Gel^®^ EX with SYBR^®^ Gold II, 2% SizeSelect pre-cast agarose gel (Invitrogen 2014, 2017). The eluted FTHFS amplicons were digested separately with restriction enzyme AluI (NEB 2020a) and Hpy188III (NEB 2020b) overnight, followed by digestion termination at 80 °C for 20 minutes. Digested amplicons were subjected to capillary electrophoresis for restriction fragment detection, which was carried out at the genotyping facility of the SNP&SEQ Technology Platform, Science for Life Laboratory, National Genomics Infrastructure, Uppsala (UGC 2018). The data from the target channel were extracted from the raw data in ABIF file format with the help of Peak Scanner™ software (Applied Biosystems 2006) and quantitative data were saved in data-frame in .csv file format. Restriction fragment data analysis was performed in Microsoft Excel 2013 (Microsoft 2013) and data visualisation was done in RStudio version 3.5.2 (RStudio Team 2015). Experimental (*Ex*) TRFs in the size range 50-640 bp were selected for clustering, quantitative analysis and visualisation. The diversity of the *Ex* oTRFs was visualised by 1) principal coordinate analysis (PCoA) using command cmdscale (package *stats* version 3.6.2) (R Core Team 2019) with Euclidean distances and 2) non-metric multidimensional scaling (NMDS) using command vegdist, metaMDS (package *vegan* version 2.5-6) (Oksanen *et al.* 2019), with Bray-Curtis dissimilarity (Bray and Curtis 1957). To fit the environmental parameters on the respective diversity plots, command envfit (package *vegan*) was used.

**Figure 1-.**
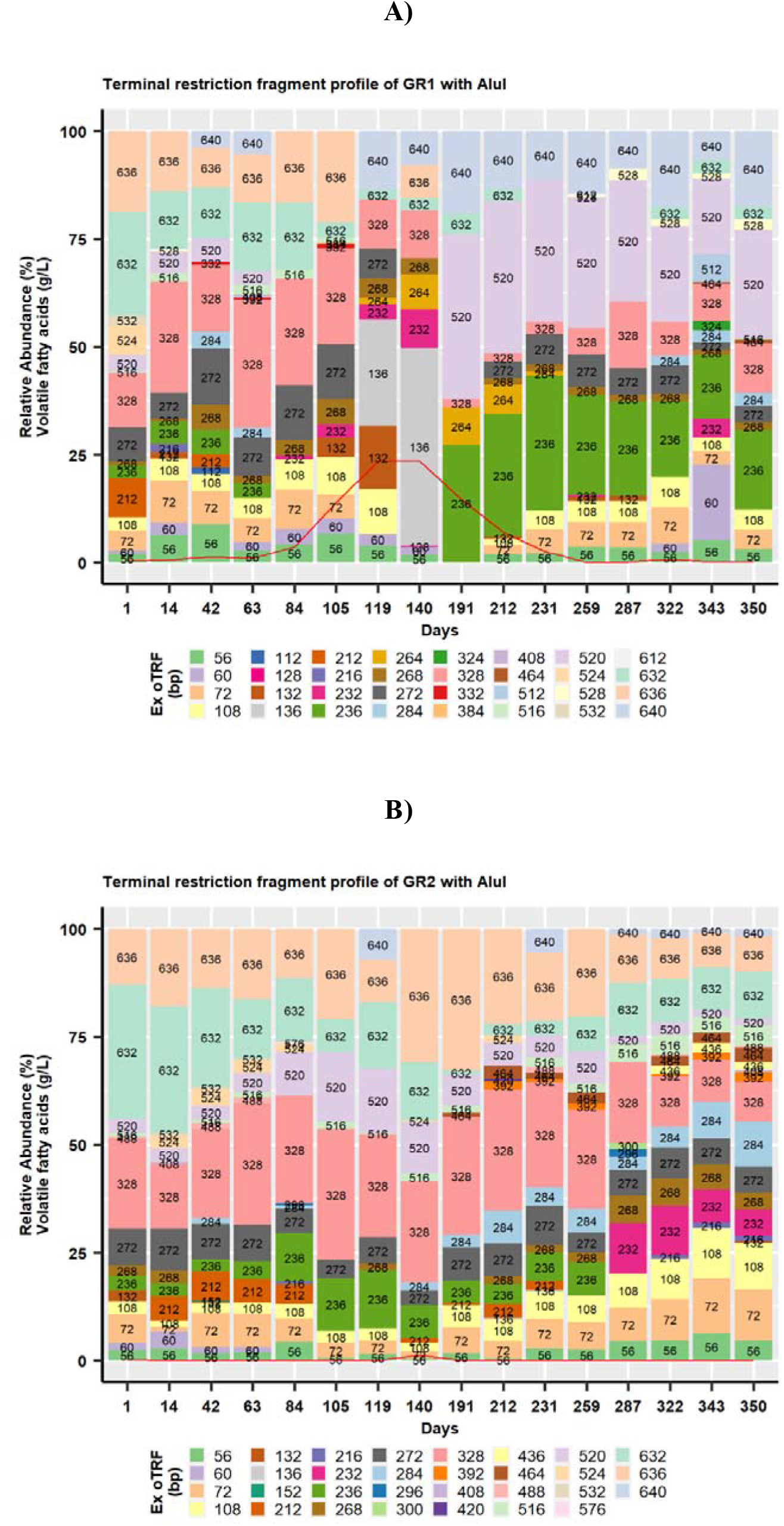
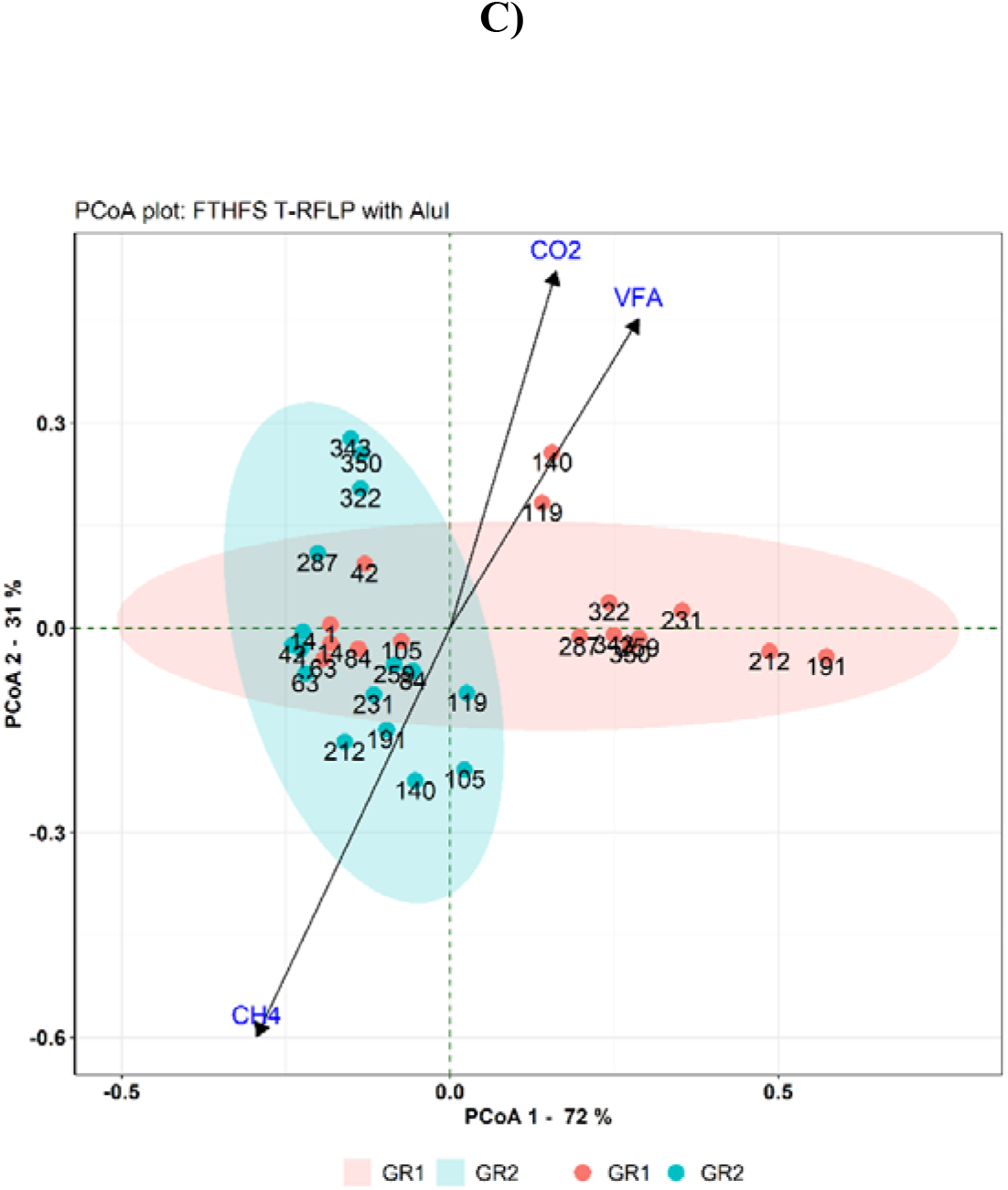
Experimental terminal restriction fragment length polymorphism (T-RFLP) profile representing *Ex* oTRF with restriction enzyme AluI for **A)** experimental reactor GR1 and **B)** control reactor GR2. The red line represents the level of total volatile fatty acids (g/L) at the respective time-point. **C)** Principal coordinate analysis (PCoA) plot showing microbial beta diversity reactors GR1 and GR2 using FTHFS T-RFLP profile with AluI. VFA, CH_4_ and CO_2_ are the environmental vectors which represent the level of total volatile fatty acids (g/L), methane content (%) and carbon dioxide content (%), respectively.

### *In silico* T-RFLP analysis for AcetoBase reference FTHFS nucleotide dataset

The T-RFLP profile of FTHFS gene fragments was simulated using the reference nucleotide dataset retrieved from AcetoBase, which consists of ~6820 non-redundant full-length taxonomically annotated FTHFS nucleotide sequences (Singh *et al.* 2019). A dataset of *in silico (IS)* PCR amplicons was generated by aligning the full-length FTHFS nucleotide reference dataset and FTHFS clone sequences generated in a previous study (Müller *et al.* 2016). Multiple sequence alignment was performed with the FAMSA alignment program (Deorowicz, Debudaj-Grabysz and Gudys 2016), with 1000 bootstrap iterations and single linkage guide tree. The resulting alignment was then trimmed to a length corresponding to the clone sequences of approximately 588 base pairs (bp). Clone sequences were removed from the alignment and all the gaps in alignment were deleted. This dataset of ungapped FTHFS nucleotide reference of ~588 bp was saved as a multi-fasta formatted file and was used as an input file for the *IS* restriction digestion and T-RFLP analysis program REDigest (Singh 2020b). For the *IS* analysis, the tagged forward (5’-CCNACNCCNNNNGGNGANGGNAA-3’; 23 bp) and reverse (5’-ATNTTNGCNAANGGNCCNCCNTG-3’; 23 bp) primer sequences were added to the input sequences. A separate *IS* analysis with the restriction enzyme AluI and Hpy188III was performed for the input file. For the quantitative and taxonomic analysis, *IS* TRFs smaller than 50 bp and greater than 640 bp (approximately [588+23+23]) were removed and excluded from the analysis. The moving average method (Smith *et al.* 2005; Fredriksson, Hermansson and Wilén 2014) was applied for binning *IS* TRFs with a size difference of ± 2 bp into an operational terminal restriction fragment unit (oTRF).

### High-throughput sequencing library preparation

The 16S rRNA gene (V3-V4) amplicon library was prepared with the primers 515F (Hugerth *et al.* 2014) and 805R (Herlemann *et al.* 2011) using the protocol described by Müller *et al.* (2016). Partial FTHFS gene amplicons were generated with the custom-indexed FTHFS primers (FWD 5’-CCNACNCCNSYNGGNGARGGNAA-3’ and REV 5’-ATNTTNGCNAANGGNCCNCCNTG-3’), for which the barcoded strategy was adopted from Hugerth *et al.* (2014). The multiplexed amplicon library was prepared by pooling equal amounts (20 ng) of each sample. For both 16S rRNA and FTHFS gene amplicon libraries, paired-end sequencing was performed (at the sequencing facility of the SNP&SEQ Technology Platform) on Illumina MiSeq with 300 base pairs (bp) read length using v3 sequencing chemistry (UGC 2018).

### Development of RibocetoBase

RibocetoBase is a subset of the 16S rRNA training dataset (Silva SSU taxonomic training data formatted for DADA2, version 138) (McLaren 2020) representing the FTHFS gene-harbouring accessions present in AcetoBase (Singh *et al.* 2019). To develop RibocetoBase, complete genomes/genomic assemblies of taxonomic identifiers for the AcetoBase accessions were downloaded from the NCBI FTP genome server (NCBI 2020) (accessed May 2020) and screened for the presence of 16S rRNA gene sequences. From among 7928 AcetoBase accessions, 6857 genomes/assemblies were successfully retrieved and 16S rRNA gene sequences were extracted with strict filtering parameters (percentage identity 95%, evalue 1e-5, window size 0) in the BLAST+ nucleotide homology search algorithm (Camacho *et al.* 2009), using the Silva SSU training dataset as reference. This showed that 1071 AcetoBase accessions were lacking complete genome/assembly sequences in the NCBI FTP genome server and could not be used for 16S rRNA gene sequence screening. All the extracted sequences were collected in a single fasta file and duplicate/redundant sequences were filtered out using the DupRemover program (Singh 2020c). RibocetoBase contains 9169 taxonomically annotated sequences for which the size distribution range is 300-2072 bp and most sequences have size around 1500 bp (Supp. Fig. S1A). The RibocetoBase database was saved in a compressed multi-fasta file format and used for acetogenic community taxonomic assignments based on 16S rRNA AmpSeq data. The length and taxonomic distribution of RibocetoBase are shown in Supp. Fig. S1B. In referring to the FTHFS gene-harbouring community inferred from 16S rRNA gene amplicons in the remainder of this paper, the term ‘Riboceto-community’ is used. Etymologically, Riboceto-community is derived from RibocetoBase and it refers to the acetogenic community inferred from 16S ribosomal RNA gene amplicons.

### High-throughput sequencing data analysis

For the 16S rRNA AmpSeq data, Illumina adapters and primer sequences from the raw data were trimmed and quality filtering for sequences with a Phred score below 20 was performed with Cutadapt (version 2.2) (Martin 2011). De-noising and generation of a taxonomy table and abundance/ASV table were done in package *dada2* (version 1.14.1) (Callahan *et al.* 2016) in R programming language (version 3.5.2)/RStudio (version 1.2.5033) (R Core Team 2013; RStudio Team 2015). Genus-level taxonomic assignment of the amplicon sequence variants (ASV) was done with the function assignTaxonomy, using the Silva taxonomic training dataset (version 138) formatted for *dada2* (Version 1) (McLaren 2020). Taxonomic assignment of Riboceto-community was done using the RibocetoBase training dataset formatted for *dada2* (this study). The results were visualised individually for the community inferred from 16S rRNA AmpSeq data (16S-community) and Riboceto-community with package *phyloseq* (version 1.30.0) (McMurdie and Holmes 2013) and vegan (version 2.5.6) (Oksanen *et al.* 2019) in RStudio (version 1.2.5033) (RStudio Team 2015).

Unsupervised FTHFS gene sequence data analysis was performed using the AcetoScan pipeline (version 1.0) (Singh *et al.* 2020). The parameters used for the AcetoScan analysis were -r 1, -m 300, -n 150, -q 21 and -c 10, while other parameters were defaults (AcetoScan user-manual). Customised visualisation of the AcetoScan results was done with package *phyloseq* (version 1.30.0) and vegan (version 2.5.6) (Oksanen *et al.* 2019) in RStudio (version 1.2.5033) (RStudio Team 2015). All data processing analyses were performed on a Debian Linux-based system with x86_64 architecture and a 3.4 GHz Intel^®^ Core™ i7-6700 processor.

## RESULTS

### Biogas reactor operation and performance profile

The performance profile of the reactors is presented in Supp. data 1. The content of carbon dioxide (%) and of methane (%) and VFA levels were used as indicators of process performance of both GR1 and GR2. In the reference reactor GR2, small fluctuations in carbon dioxide and methane content were observed, with no accumulation of organic acids. With the increased organic load in GR1 (Δ ~1.59 g VS L^-1^ day^-1^), an increase in carbon dioxide content and a decrease in methane content were detected, indicating deteriorating performance of GR1. An increase in the VFA concentration was also noticed in GR1 from day 84, with a peak of 24.6 g/L on day 133. On returning the OLR for GR1 to ~2.5 g VS L^-1^ day^-1^ (~day 125), an increase in methane content was recorded, followed by a decrease in the concentration of total VFA and carbon dioxide content (Supp. Data 1, Supp. Fig 2). The increase in methane content and decrease in carbon dioxide content indicated recovery of the process performance to a level similar to GR2.

### Quantitative analysis of terminal restriction fragments

Quantitative analysis of *Ex* TRFs was done using the experimental data stored in data-frames during the preliminary analysis. *Ex* TRFs differing by ± 2 bp in size were binned to produce an *Ex* oTRF. This binning of *Ex* TRFs helped to identify, quantify and visualise the *Ex* TRFs (Fig. 1, Supp. Fig. S3). Similarly, *IS* TRFs were binned in *IS* oTRFs to compare with *Ex* oTRFs. However, the binning strategy caused differences in the size of a TRF generated from *IS* restriction digestion of a sequence and an oTRF from the same sequence (Fig. 2). Therefore, in the absence of an *IS* TRF equal to an *Ex* oTRF, the taxonomy of the *IS* oTRF (± 2 bp) is considered in the discussion. The *Ex* oTRFs 636 and 640 bp were the unrestricted fragments, and are thus not considered in the discussion.

**Figure 2-.**
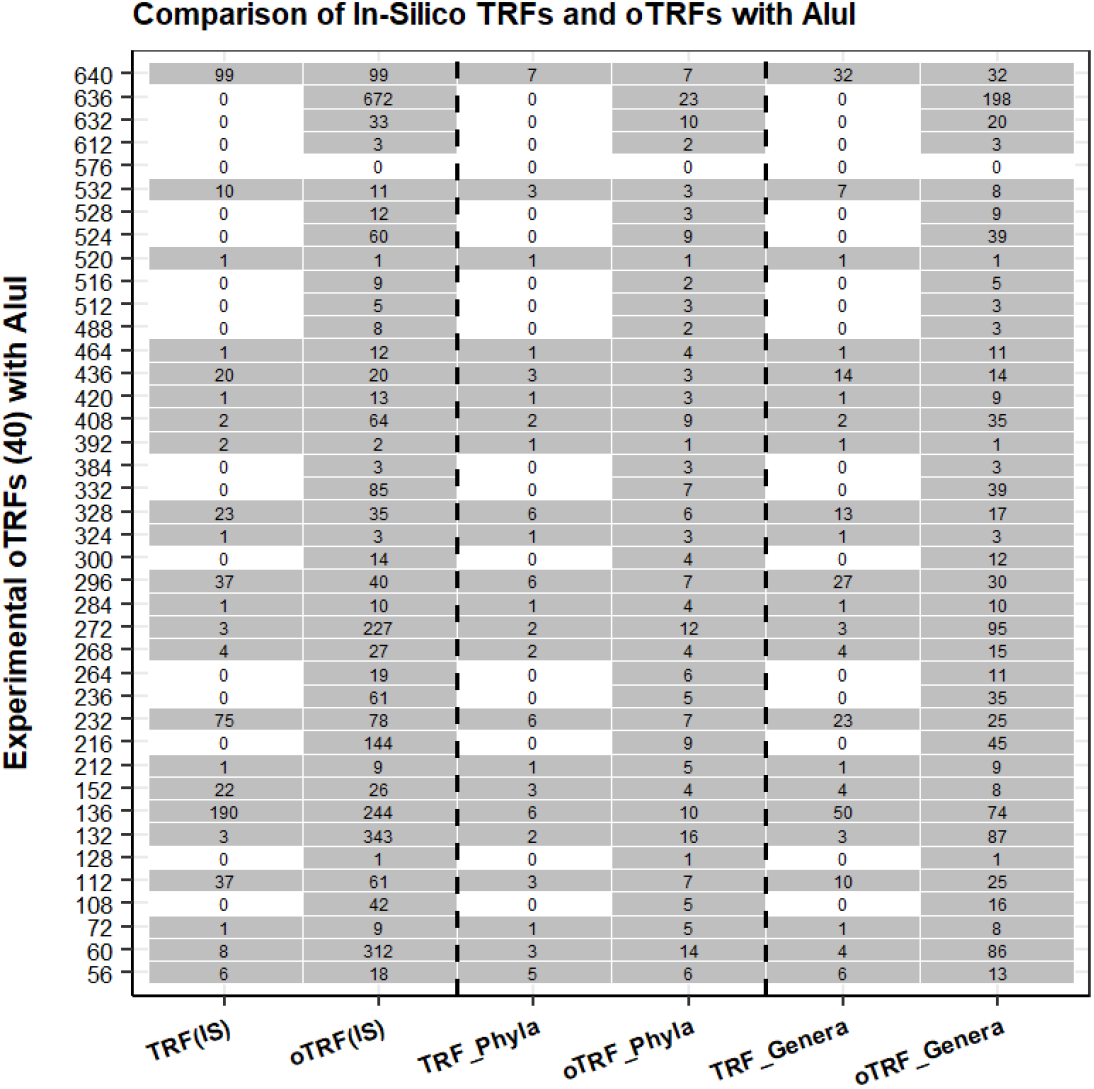
Tabular plot representing the T-RFLP profile during comparison of experimental oTRF versus *in silico* TRF, *in silico* oTRF and count of taxa (phyla and genera) with restriction enzyme AluI.

### Experimental T-RFLP profiles generated from GR1 and GR2

The restriction profile of FTHFS gene fragments from GR1 revealed significantly different community dynamics during (1-125 days) and after (126-350 days) the disturbance phase. The community composition in GR1 during the initial phase of disturbance was very similar to that in GR2, but gradually changed under the influence of disturbance and increasing total VFA concentration. The restriction profile from GR1 and GR2 with AluI consisted of 32 and 31 *Ex* oTRFs, respectively, with total 40 unique *Ex* oTRFs. Of these 40 *Ex* oTRFs, 23 were seen in both reactors and 17 *Ex* oTRFs were unique to either GR1 (nine unique *Ex* oTRFs, of size 112, 128, 264, 324, 332, 384, 512, 528 and 612 bp) or GR2 (eight unique *Ex* oTRFs, of 152, 296, 300, 392, 420, 436, 488 and 576 bp). Among the unique *Ex* oTRFs in GR1, two (264 and 528 bp) were observed to have relative abundance (RA) >1%, while none of the eight unique *Ex* oTRFs in GR2 was seen to have RA >1% during the experimental period (Fig. 1). In GR1, the disappearance of *Ex* oTRFs 236 and 520 bp during the early disturbance phase (day 63) and their reappearance during the recovery phase (day 191) were strongly connected with the increasing/high and decreasing/low level of total VFA, respectively. *Ex* oTRFs 132, 136 and 264 bp were only observed between day 105 and 231, whereas *Ex* oTRF 72 bp disappeared between day 119 and 191, when VFA accumulation was high. *Ex* oTRF 528 bp was only observed (RA >1%) from day 259 to 350 (Fig. 1A). *Ex* oTRFs 272, 328 and 632 bp were the most prominent *Ex* oTRFs before the VFA levels increased. No such significant changes were noticed in GR2, except that *Ex* oTRF 232 bp disappeared and *Ex* oTRF 236 bp appeared between day 287 and day 350 (Fig. 1B). Principal coordinate analysis (PCoA) of the AluI restriction profile resulted in close clustering of samples from GR2, while samples from GR1 were dispersed under the influence of increased carbon dioxide content (%) and total VFA concentration (g/L). The environmental vector for methane content (%) was opposite to that for carbon dioxide content and total VFA concentration, indicating that methane content decreased when the carbon dioxide content and total VFA concentration increased and *vice versa* (Fig. 1C). Similar to the AluI restriction profiles of GR1 and GR2, the restriction profile with restriction enzyme Hpy188III also resulted in visually different *Ex* oTRF composition and dynamics in GR1 compared with GR2. The *Ex* oTRF profile in GR1 was similar to that in GR2 during the initial phase of disturbance, but gradually changed with the increase, followed by a decrease, in total VFA concentration. The experimental T-RFLP profile generated from Hpy188III restriction digestion is presented in additional text (Fig. A1). Results of NMDS analysis for both GR1 and GR2 with restriction enzyme AluI and Hpy188III showed similar trends to those seen in PCoA analysis (Supp. Fig. S7A and S7B).

### *In silico* T-RFs from AcetoBase FTHFS nucleotide dataset

The *IS* restriction digestion of the FTHFS nucleotide dataset with restriction enzyme AluI resulted in 360 *IS* TRFs, among which 326 *IS* TRFs were within the size range 50-640 bp (Supp. Fig. S3A). With restriction enzyme Hpy188III, the total number of *IS* TRFs was 399, of which 363 *IS* TRFs were within the size range 50-640 bp (Supp. Fig. S3B). These *IS* TRFs with size ranging between 50 and 640 bp were used for the oTRF binning. After binning the *IS* TRFs, 140 and 142 oTRFs were generated for AluI and Hpy188III, respectively, and used for further analysis and comparison (Fig. 2, Supp. data 2, Supp. Fig. S4).

### Comparison of experimental and *in silico* T-RFLP profile for AluI and taxonomic prediction of TRFs

The taxonomic predictions for the TRFs were made by comparing the *Ex* oTRF profile to the *IS* TRF profile generated from the sequences of known taxonomy. The T-RFLP profiles of *Ex* oTRFs from GR1 and GR2 generated with AluI were different from those obtained for the *IS* TRFs in terms of the size and number of the restriction fragments (Fig. 2, Supp. Data 2). The taxonomy of the *Ex* oTRF 72 bp represented only one hit in the *IS* analysis, belonging to the phylum Actinobacteria and genus *Salinibacterium*, while *IS* oTRF represented nine hits (70 (2), 71 (2), 72 (1) and 73 (4) bp) belonging to five phyla and eight genera. *Ex* oTRF 132 bp was represented by three *IS* TRFs (belonging to two phyla and three genera) and by 343 *IS* oTRFs (16 phyla and 87 genera). The most dominant *Ex* oTRF, *i.e.*, 136 bp, between day 119 and 140 (Fig. 1A) represented 190 and 244 hits in the *IS* TRF and *IS* oTRF profile, respectively (Fig 2). The *IS* TRF hits belonged to six phyla and 50 genera and the *IS* oTRF hits belonged to 10 phyla and 74 genera (Fig. 2, Supp. Data 2), including *Clostridium* (*C. ultunense, C. beijerinckii, C. perfringens, C. formicaceticum*), *Clostridioides, Dorea, Eubacterium, Prevotella, Proteus, Sporomusa* and *Terrisporobacter etc*.

*Ex* oTRF 264 bp, which was unique to GR1, did not appear in *IS* restriction digestion, but *IS* TRFs 262 (7), 263 (11), 265 (1) and 266 bp (5) were generated. This can be explained by the fact that experimental and *IS* TRFs were clustered (±2 bp), and thus specific *IS* TRF of 264 bp may not exist. However, if taxonomy of *IS* TRFs generated from the reference dataset is considered, six out of seven *IS* TRFs of size 262 bp belonged to the genus *Treponema. IS* TRF 263 bp was related to the genus *Blautia* and *IS* TRF 265 bp was generated from the genus *Moorella* (Supp. Data 2). One out of five *IS* TRFs of size 266 bp was from the genus *Acetobacterium. Ex* TRF 236 bp was not generated in *IS* analysis, but clustering produced 61 *IS* oTRFs of size 236 bp, which belonged to five phyla and 35 genera (Fig. 2, Supp. Data 2). The *IS* oTRF of 236 bp consisted of *IS* TRF 235 (60) and 237 (1) bp and were taxonomically associated with genera like *Blautia, Clostridium, Hungateiclostridium, Oxobacter, Prevotella. Ex* oTRF 520 bp, present in high relative abundance in GR1, represented only one hit in the *IS* analysis, belonging to the phylum Fusobacteria. However, when *IS* TRF of size 522 bp was also considered, there were 60 *IS* TRFs clustered into *IS* oTRF of 524 bp. Among these *IS* oTRF, taxonomically nine phyla and 39 genera were represented by *IS* oTRF 524 bp, including species (Candidatus) *Cloacimonetes* bacterium HGW-Cloacimonetes-1. The *IS* TRF profile was lacking *Ex* oTRFs of 632 bp. However, 33 *IS* oTRFs of 632 bp were generated, belonging to 10 phyla and 20 genera. The taxonomy predicted for the major AluI *Ex* oTRFs and their relative abundance is presented in Supp. Fig. S5. *Clostridium* was the most abundant and diverse genus, followed by *Eubacterium*, represented by the major *Ex* oTRFs generated by AluI.

### Riboceto-community structure and dynamics

Analysis of Riboceto-community revealed three known phyla and one unknown bacterial phylum (Bacteria.NA) with RA >1%. These phyla were Actinobacteriota (RA 1-5%), Firmicutes (6-52%), Bacteriodota (28-90%) and Bacteria.NA (1.1-50.6%) in both GR1 and GR2 (Fig. 3A). At the class level, eight classes were found to have RA >1%, of which five major classes were Bacilli (RA 1-18%), Bacteroidota (38-90%), Bacteria.NA (1.2-51%), Clostridia (1.34-32%) and an unknown class of Firmicutes (1.2-13.5%) in both GR1 and GR2. At the family level, the families that differed most between the experimental and control reactors during the disturbance phase in G1 were an unknown family of Bacteroidales and Tannerellaceae (Supp. Fig. S6). The RA of Bacteroidales.NA increased and decreased with the rising and falling level of total VFA in GR1, respectively. Similarly, the RA of Tannerellaceae decreased and increased with the change in VFA level of GR1. The family Bacteria.NA in GR1 disappeared and reappeared with the rise and fall in total VFA level. In contrast, these unknown families of Bacteroidales and Tannerellaceae had a relatively stable presence throughout the operational phase in GR2. The unknown bacterial family Bacteria.NA was observed to have fluctuating RA in GR2. The family Hungateiclostridiaceae was only observed in GR1 (RA >2%) (Suppl. Fig. S6). Genus-level community analysis (RA >1%) revealed patterns similar to the family-level analysis for Bacteria.NA and Clostridia.NA genera (Fig. 3B). The most noteworthy change was the appearance of the genera *Tissierella* (day 84-133) and *Proteiniphilum* (day 95-140), and an unknown member of the family Tannerellaceae (day 105-182), with the increase in total VFA. These genera were observed sporadically in GR2, but RA was found not to be above 2.2% during the whole operational phase. PCoA analysis with weighted UniFrac distance matrix indicated distinct clustering of the samples from GR1 and GR2, and the environmental vectors carbon dioxide content (%) and total VFA (g/L) were opposite to methane content (%) (Fig. 3C). The NMDS analysis of community based on 16S rRNA gene sequences represented the separate clustering of the samples from GR1 and GR2 and dispersal of the samples under the influence of environmental vectors (CH_4_, CO_2_ and VFA) (Suppl. Fig. S7C).

**Figure 3-.**
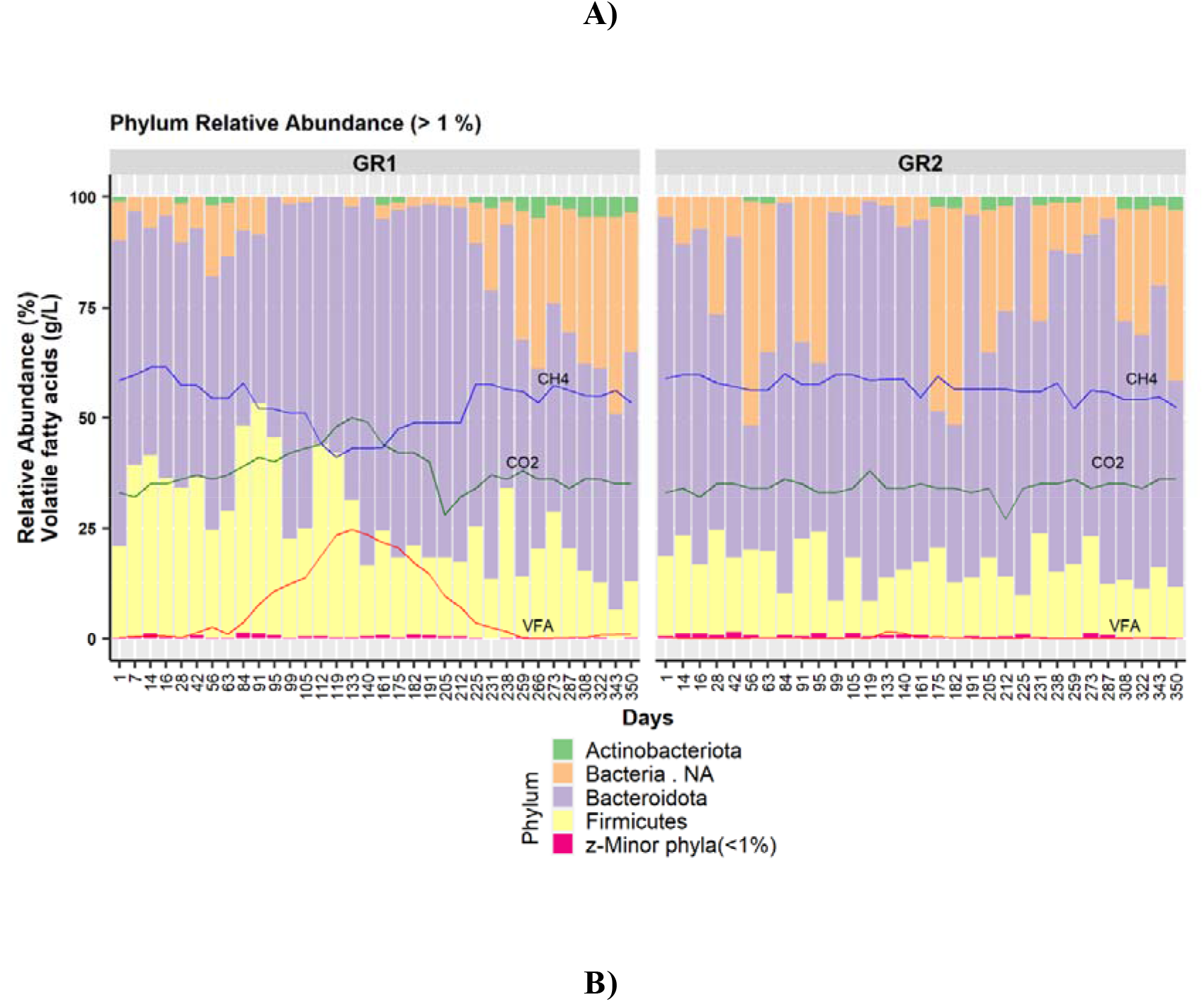
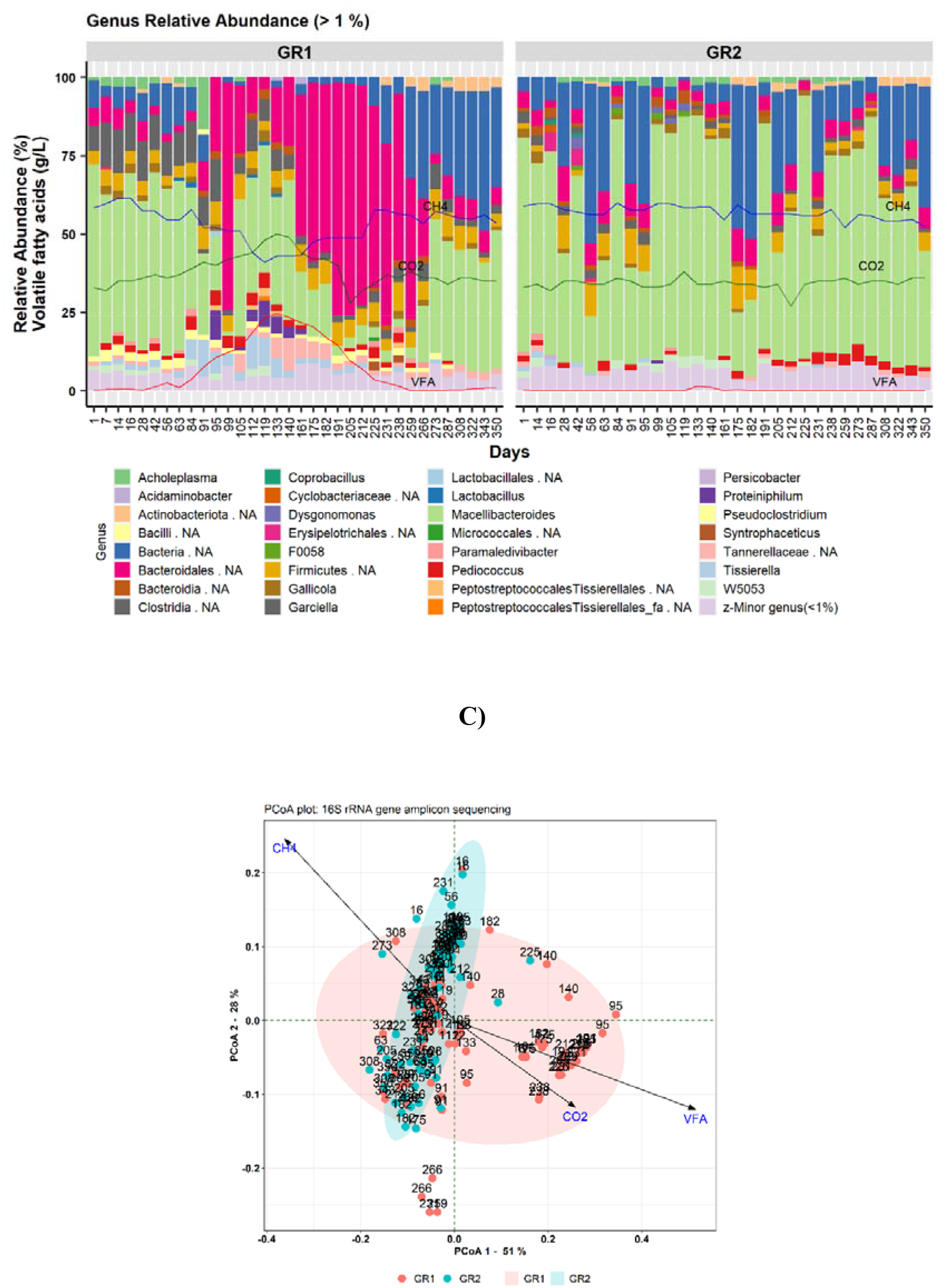
Bar plot representing Riboceto-community in experimental reactor GR1 and control reactor GR2 at the **A)** phylum level (relative abundance (RA) >1%) and B) genus level (RA >1%). VFA, CH_4_ and CO_2_ represent the level of total volatile fatty acids (g/L), methane content (%) and carbon dioxide content (%), respectively. **C)** Principal coordinate analysis (PCoA) plot with weighted UniFrac distance matrix visualising beta diversity of the whole microbial community in samples from reactors GR1 and GR2 under the influence of the environmental parameters carbon dioxide content (%), total VFA (g/L) and methane content (%).

### Potential acetogenic community structure based on FTHFS gene amplicons – “FTHFS-community”

The high-throughput sequencing followed by data analysis with AcetoScan for FTHFS gene amplicons indicated that, at the phylum level, six phyla (Actinobacteria, Candidatus Cloacimonetes, Firmicutes, Proteobacteria, Spirochaetes and Synergistetes) had RA >1%. Candidatus Cloacimonetes and Firmicutes were the most abundant phyla, making up approximately 80% of the total community in both GR1 and GR2 during the study period. In GR1, phylum Candidatus Cloacimonetes showed increased RA during days 1-125 and decreased RA during days 125-350. The phyla Candidatus Cloacimonetes and Firmicutes were seen to be relatively stable (with smaller occasional fluctuations in RA) in GR2 (Fig. 4A). At the class level, community structure and dynamics seen for the class Actinobacteria, unclassified Candidatus Cloacimonetes and Clostridia were similar to those seen at the phylum level for GR1 and GR2. However, an increase in RA >1% for the class Negativicutes (1.2-3.2 %) was noted in GR1 from around day 91 to day 182, in line with the increase in VFA levels. Occasional appearance (RA >1%) of the classes Tissierellia and Spirochaetia was also seen, but only in GR1. At the order level, RA of the orders Selenomonadales, Tissierelliales and Spirochaetales were found to increase (up to 8%) and decrease (to ≤1%) with the increase and decrease in total VFA levels in GR1, respectively. These orders were not observed in GR2, where ~90% of the community was composed mainly of an unknown order of phylum Candidatus Cloacimonetes. At the genus level, appearance and increase (3-40%) in the RA of an unclassified Peptococcaceae genus with the increase in VFA levels was also observed, followed by appearance of the genus *Eubacterium* (RA 5-11%) from day 99 to day 238. The genus *Marvinbryantia* also showed increasing RA (7-40%) from day 133 to day 82, and then declined again to RA <1%. The genera *Eubacterium* and *Marvinbryantia* were not observed in RA >1% and unclassified Peptococcaceae did not exceed RA >2% in control reactor GR2 (Fig. 4B).

**Figure 4-.**
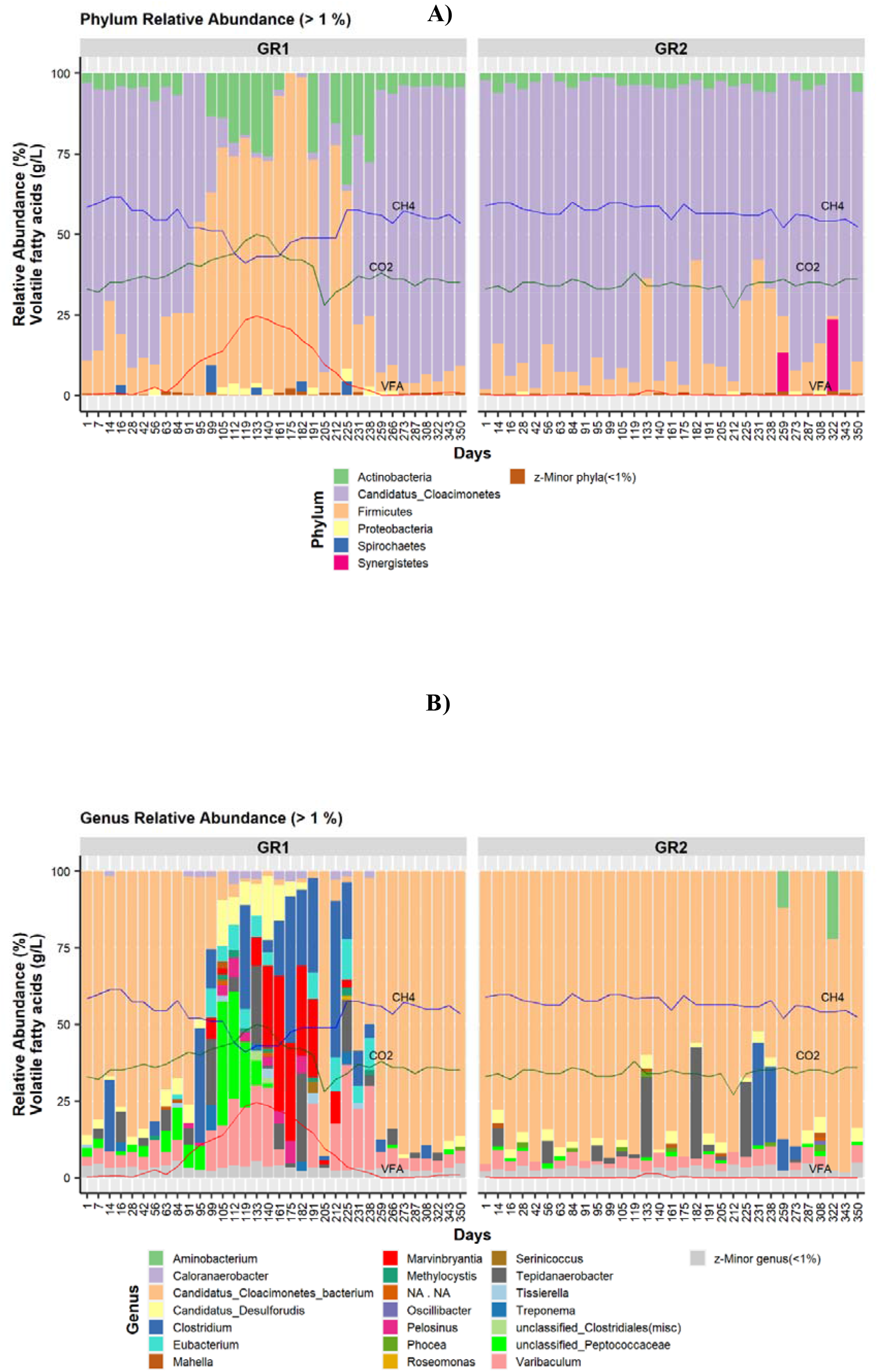
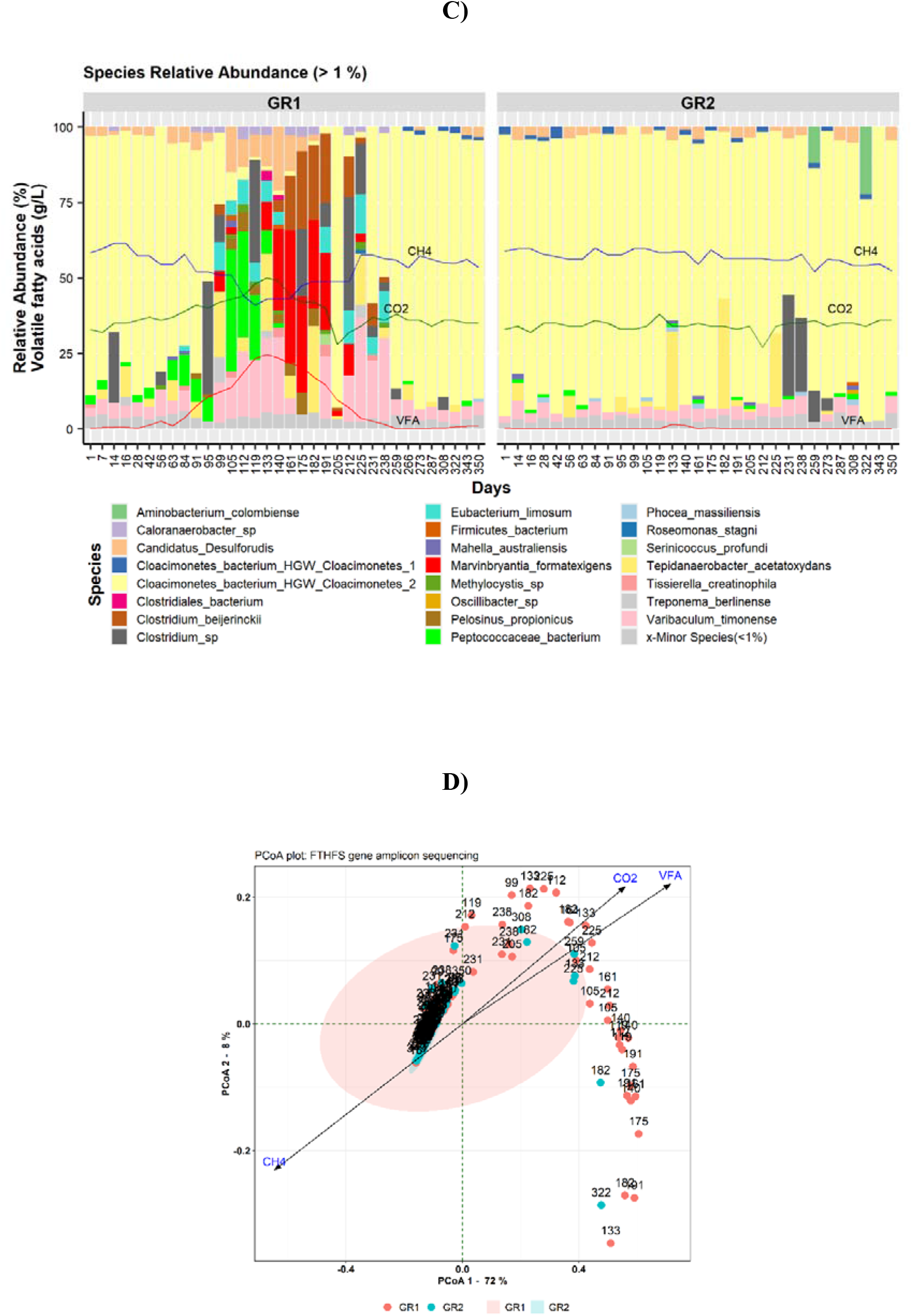
Bar plot of acetogenic community based on FTHFS gene amplicons in experimental reactor GR1 and control reactor GR2 at **A)** phylum level (relative abundance (RA) >1%), **B)** genus level (RA >1%) and **C)** species level (RA >1%). VFA, CH_4_ and CO_2_ represent the level of total volatile fatty acids (g/L), methane content (%) and carbon dioxide content (%), respectively. **D)** Principal coordinate analysis (PCoA) plot with weighted UniFrac distance matrix visualising beta diversity of the potential acetogenic community inferred from FTHFS gene sequencing in samples from reactors GR1 and GR2 under the influence of environmental parameters carbon dioxide content (%), total VFA (g/L) and methane content (%).

At the species level, (Candidatus) *Cloacimonetes* bacterium HGW-Cloacimonetes-1 was only seen to have RA >1% after day 259 in GR1, while in GR2 it was seen (RA >1-3%) throughout the operational phase. Moreover, the species *Eubacterium limosum, Clostridium beijerinckii, Marvinbryantia formatexigens, Treponema berlinense, Tissierella creatinophila, Caloranaerobacter* sp. and *Pelosinus propionicus* were only detected in GR1 during the disturbance phase or when the level of total VFA was high. The unknown genus of the family Peptoccocaceae was observed to have increasing RA (1-40%) in GR1 with the increase in total VFA (0.5-18.6 g/L) and its RA decreased (to ≤1%) with the decrease in total VFA level, while it was detected to have almost constant RA of ≤2% in GR2 (Fig. 4C). PCoA analysis of the potential acetogenic community inferred from FTHFS AmpSeq indicated very tight clustering of the samples from the control reactor G2 (Fig. 4D). The samples from the experimental reactor GR1 were dispersed along the environmental vectors for carbon dioxide content (%) and total VFA (g/L), and opposite to methane content (%) (Fig. 4D). NMDS analysis of FTHFS-community indicated trends similar to 16S-community beta diversity, where GR1 and GR2 samples showed distinct clusters based on the experimental and recovery period in GR1 and influence of environmental vectors (CH_4_, CO_2_, VFA) (Suppl. Fig. S7D).

## DISCUSSION

All the methods used in this study for community analysis revealed a similar pattern in terms of community dynamics during the disturbance and recovery phase in the experimental reactor (GR1) and relatively stable community structure in the control reactor (GR2). An increase in carbon dioxide content (%) and total VFA concentration (g/L) and decrease in methane content (%) were the main indicators of disturbance in both FTHFS- and 16S-community structure for GR1 compared with GR2. However, there were considerable differences in the community profile when taxonomy associated with the method taken into consideration. Differences arose because different methods produce slightly differing results and have their advantages and disadvantages. However, detailed comparison of the methods revealed 1) similarity in temporal dynamics, taxonomy and RA abundance of respective taxa/TRFs and 2) some differences and plausible reasons for these.

### Comparison of T-RFLP profile and structure and dynamics of Riboceto-, 16S-, FTHFS-community

#### FTHFS gene-based community dynamics: T-RFLP versus AmpSeq

On comparing the FTHFS gene-based community and taxonomic predictions of AluI *Ex* oTRFs, it was observed that unknown bacteria of the family Peptococcaceae and *M. formatexigens* were present in significantly high relative abundance in FTHFS-community in GR1, but were not detected in the *Ex* oTRF profile (via its taxonomic prediction). This was likely because, as the *in silico* digestion results indicated, the TRFs generated by these two species were smaller than 50 bp (Supp. Data 2), and thus were not included in the T-RFLP analysis. In the *IS* T-RFLP profile of AluI and Hpy188III, some *IS* oTRFs were represented by several genera, such as *Clostridium* (*C. ultunense, C. beijerinckii, C. perfringens, C. formicaceticum*), *Clostridioides, Dorea, Eubacterium, Prevotella, Proteus* and *Sporomusa, Terrisporobacter etc.* Although no exact match for a particular genus (species) was found for *IS* oTRFs, all of these genera are relevant for the community dynamics because they include most of the known acetogens (Drake, Gößner and Daniel 2008; Singh *et al.* 2019). The appearance and increase in RA of certain *EX* oTRFs in GR1 and their predicted taxonomy illustrate the important role of these acetogens in VFA metabolism. Comparison of dynamics deduced from the FTHFS gene T-RFLP and AmpSeq indicated similar trends in the RA of associated or predicted taxa. However, our *in silico* analysis suggested that accurate identification and prediction of exact taxa based on TRF identity is not feasible.

#### Congruence of RibocetoBase taxonomic annotations with 16S-community

Analysis of Riboceto-community (Fig. 3) showed similar trends to those seen for 16S-community dynamics (additional text, Fig. A2). At phylum level, the major phyla observed in Riboceto-community and 16S-community were very similar, although increased resolution for these dynamics was observed for Riboceto-community (*e.g.*, acetogenic community) (Fig. 3A) compared with 16S-community (additional text, Fig. A2A). At phylum level, 16S-community in GR1 showed complete disappearance of the phylum Cloacimonadota between day 95 and day 225 and presence of phylum Actinobacteriota in only a few samples. In line with this, Riboceto-community indicated reduced RA of the phylum Cloacimonadota during days 95-225, although unlike in 16S-community it did not completely disappear. In Riboceto-community, the phylum Actinobacteriota (RA >1% intermittently) was observed in both GR1 and GR2. 16S-community showed similar trends to Riboceto-community at phylum level, where Candidatus Cloacimonetes was highly reduced during the disturbance but did not disappear and Actinobacteriota was observed in both reactors. In the comparison of Riboceto-community (Fig. 3A) and 16S-community (additional text, Fig. A2A), it was noticed that the phylum Cloacimonadota was annotated “NA” in Riboceto-community. This is because RibocetoBase taxonomy is based on NCBI taxonomy (Federhen 2012), while 16S rRNA gene reference dataset taxonomy is based on Arb-Silva (Quast *et al.* 2013), recently amended with GTDB taxonomy (Chaumeil *et al.* 2019; Parks *et al.* 2020). If the taxonomy of the “NA” sequences in the Riboceto-community fraction is traced using Silva aligner (Quast *et al.* 2013) or RDP sequence match (Cole *et al.* 2014), and taxonomy is compared to GTDB taxonomy, the taxonomic affiliations of ASVs can be compared/validated for “NA” taxa. Thus, these differences in the taxonomic classification systems created differences between the taxonomy associated with the community dynamics. Other reasons for the assignment of phylum Cloacimonadota (GTDB taxonomy) or Candidatus Cloacimonetes (NCBI taxonomy) as unknown bacterial phylum in the RibocetoBase dataset are: 1) lack of complete genomes/genomic assemblies for the Candidatus Cloacimonetes accessions present in AcetoBase and 2) absence of 16S rRNA gene sequences in these genomes/genomic assemblies, with 5S or 23S subunits mostly present in these genomic assemblies. Thus, the 16S rRNA gene sequence could not be extracted from the Cloacimonadota/Candidatus Cloacimonetes genomes for the RibocetoBase dataset and identified ASVs could not be annotated. A comparison of FTHFS-community and 16S-community also indicated the reason for the high RA of Actinobacteria in the FTHFS profile and its low RA in the 16S rRNA gene profile. The genus *Varibaculum timonense*, which was previously classified in the family Actinomycetaceae (AcetoBase) (Singh *et al.* 2019) under old NCBI taxonomy (Federhen 2012) has been reclassified to *Urmitella timonensis* in the family Tissierellaceae (GTDB 2020a) according to GTDB taxonomy (Chaumeil *et al.* 2019). Hence, amendments in RibocetoBase taxonomy according to GTDB taxonomy would be required for correct taxonomic annotations and community profiling. However, indirect inference using RibocetoBase, *i.e.* FTHFS gene-harbouring bacteria from AcetoBase, would be a satisfactory method for potential acetogenic community profiling in cases where the 16S rRNA gene can be used as the marker of choice.

#### Resolution of acetogenic community structure: functional gene versus taxonomic marker

The FTHFS gene is a functional gene marker of the acetogenic community. A prominent genus in FTHFS-community *i.e.*, Peptococcaceae bacterium (Fig. 4B), was not observed in 16S-community (additional text, Fig. A2B). Further analysis of the Peptococcaceae bacterium operational taxonomic unit (OTU) sequence showed that this OTU was 88.7% similar to the species Peptococcaceae bacterium 1109. There are several possible reasons why Peptococcaceae bacterium 1109 could not be detected in 16S-community, *e.g.* 1) it has been reclassified as class Limnochordia, family DTU010 and genus 1109 according to recent GTDB taxonomy (GTDB 2020b), 2) this species was not targeted by our 16S rRNA gene primers (515F-805R). The genera H1, PeH15, *Proteiniphilum* and LNR_A2-18, which were predominantly seen in 16S-community, were not observed in FTHFS-community. This was because H1, PeH15 and *Proteiniphilum* belong to the phylum Bacteroidota and LNR_A2-18 belongs to the Cloacimonadota. As Bacteroidota do not include any known acetogen (Drake, Gößner and Daniel 2008; Pierce *et al.* 2008; Müller and Frerichs 2013; Singh *et al.* 2019), our FTHFS primers might be unable to target these genera. The genus LNR_A2-18 is not included in AcetoBase because there is no information available for this genus apart from the 16S rRNA gene sequence.

*Marvinbryantia formatexigens* was represented in high RA in FTHFS-community during the disturbance period in GR1 (Fig. 4C). This species is a known acetogen (Drake, Gößner and Daniel 2008; Müller and Frerichs 2013; Singh *et al.* 2019) belonging to the family Lachnospiraceae. However, neither *M. formatexigens* nor family Lachnospiraceae was detected to have a significant presence in 16S-community during the disturbance period in GR1 (Fig. A2, Supp. Fig. S8). To validate the coverage of bacterial community by our FTHFS primers, the top phyla and classes from FTHFS-community and 16S-community were compared. In the comparison at phylum level, 16S-community showed higher coverage of Firmicutes than in FTHFS-community (Supp. Fig. S9A, S9B). This is reasonable, since not all Firmicutes contain the FTHFS gene and were therefore targeted in 16S-community, but FTHFS-community. At class level, FTHFS-community illustrated better coverage of class Negativicutes compared with 16S-community (Supp. Fig. S9C, S9D). Since phylum Cloacimonadota has been proposed as an indicator taxon pertaining to reactor disturbances (Calusinska *et al.* 2018; Klang *et al.* 2019; Poirier *et al.* 2020), its correct profiling is very important. Fluctuations (near-disappearance) in RA of the phylum Cloacimonadota and class Cloacimonadia were observed in the control reactor profile of 16S-community (Supp. Fig. S9B, S9D) but not that of FTHFS-community (Supp. Fig. S9A, S9C). Thus, comparative analysis of FTHFS-community and 16S-community illustrated that our FTHFS gene-based sequencing approach appears more accurate and reliable in targeting the correct coverage, dynamics and classification of known and potential acetogens.

#### Acetogenic community dynamics in the biogas environment

Since there is a lack of studies targeting acetogenic bacteria and examining their role in biogas digester environments, a clear causal attribution for the microbial community changes regarding process parameters is challenging. However, many studies have reported high importance of the acetogenic community in biogas environments and the present study contributed further insights regarding the acetogenic community structure in AD environments. This study also clearly revealed the dynamics and changes in the structure of this community with an increase and decrease in total VFA concentration. Among the major taxa detected in FTHFS-community, the dynamics of the phylum Candidatus Cloacimonetes, genus Peptococcaceae bacterium and *Marvinbryantia* are worth mentioning. Candidatus Cloacimonetes was found to be highly reduced during the disturbance phase. It has previously been reported to be present in high abundance in high-ammonia systems and is suggested to have a specialist function (syntrophic propionate-oxidation) (Müller *et al.* 2016; Poirier *et al.* 2020) and to be a potential acetogenic candidate (Pelletier *et al.* 2008; Juste-Poinapen *et al.* 2015; Lucas *et al.* 2015; Nobu *et al.* 2015; Ahlert *et al.* 2016; Stolze *et al.* 2018, 2016; Calusinska *et al.* 2018; Nazina *et al.* 2018; Braz *et al.* 2019; Klang *et al.* 2019). In line with the results in the present study, a decrease abundance in the phylum Cloacimonetes in association with process disturbance has been seen previously (Klang *et al.* 2019). The role of the genus Peptococcaceae bacterium or the family Peptococcaceae has not yet been well studied, but several studies suggest a role as an acetogen/syntrophic bacterium in natural and methanogenic environments (Müller *et al.* 2010, 2016; Liu and Conrad 2011; Kato and Yumoto 2015; Tveit *et al.* 2015; Liu *et al.* 2016). Peptococcaceae bacterium 1109 is suggested to be a syntrophic acetate oxidising bacterium (Buettner *et al.* 2019). Therefore the increased RA of this species during the VFA increase in the present study could potentially be associated with high levels of acetate in the disturbance phase, which are suggested to be of importance for other known syntrophic acetate-oxidising bacteria (Westerholm, Moestedt and Schnürer 2016). *Marvinbryantia formatexigens*, which showed increased RA during the period of VFA decrease, is a well-known acetogen in the gut environment (Wolin *et al.* 2003; Rey *et al.* 2010), but its role in biogas environments is not well studied. However, the species has cellulolytic and saccharolytic activity and produces acetate as a sole or main product from its metabolism (Wolin *et al.* 2003). In summary, in this study we successfully revealed the temporal dynamics of the acetogenic community and changes in the structure of this community structure due to perturbation.

## CONCLUSIONS

This study compared temporal changes in the acetogenic community in biogas reactors under the influence of induced disturbance, using three methods widely applied for microbial analysis. Despite differences and limitations of the 16S rRNA gene AmpSeq, T-RFLP and AmpSeq of FTHFS gene methods, all were able to track the temporal dynamics of the microbial community and coherence was found in community profiles at higher taxonomic levels inferred by individual methods. However, T-RFLP and AmpSeq of FTHFS gene were found to be more descriptive and reliable in tracking the dynamics of the acetogenic community. Overall, high-throughput FTHFS gene sequencing of barcoded samples and unsupervised analysis with AcetoScan was found to be a more promising method for monitoring acetogenic community dynamics in biogas reactors than 16S rRNA gene sequencing targeting the whole bacterial community and laborious/limited T-RFLP community profiling. If the recent taxonomic changes and differences between Arb-Silva/GTDB and AcetoBase/NCBI taxonomy are masked, AmpSeq of FTHFS gene may accurately reveal the microbial profile and dynamics of the acetogenic community in anaerobic digesters and in various natural environments (soil, marine, lake sediments, hot springs *etc.*) and gut/oral (insects, animals, human) environments.

## FUTURE PERSPECTIVES

The FTHFS gene has been used for more than three decades to identify/track the dynamics of potential acetogenic candidates. In this study, we developed an alternative approach for the same purpose, using 16S rRNA gene sequences from FTHFS gene-harbouring bacteria. This approach can have wide application in tracking the microbial population when complemented with indirect analysis of potential acetogenic candidates. Further development and improvement of RibocetoBase with more acetogen-specific 16S rRNA gene sequences and amending the NCBI taxonomy with GTDB taxonomy will further assist in identification of FTHFS gene-harbouring candidates with acetogenic potential. Use of FTHFS gene AmpSeq in screening diverse environments could also provide a deeper understanding of acetogenic community structure and its temporal dynamics in natural or constructed environments.

## SUPPLEMENTARY DATA

Supplementary data are available online.

## DATA AVAILABILITY STATEMENT

The raw sequence data from the multiplexed Illumina MiSeq sequencing for the FTHFS and 16S rRNA gene have been submitted to NCBI, with BioProject accession number PRJNA687725 and PRJNA687735, respectively.

## ACKNOWLEDGEMENTS

We would like to thank Bioprocess Engineer Simon Isaksson for his help in reactor operation, sampling and data collection.

