## Additional text for "Profiling acetogenic community dynamics in anaerobic digesters - comparative analyses using next-generation sequencing and T-RFLP"

Running title: Comparative analysis of acetogenic community

Abhijeet Singh^1*^, Bettina Müller^1^, Anna Schnürer^1*^

^1^Anaerobic Microbiology and Biotechnology Group, Department of Molecular Sciences, Swedish University of Agricultural Sciences, Almas Allé 5, Uppsala, SE-750 07, Uppsala, Sweden

### **Experimental Hpy188III T-RFLP profiles generated from GR1 and GR2**

The restriction digestion of partial FTHFS gene amplicons with restriction enzyme Hpy188III resulted in 33 unique *Ex* oTRFs. The numbers of *Ex* oTRFs observed in GR1 and GR2 were 29 and 22 respectively. Out of 33 unique *Ex* oTRFs, 17 *Ex* oTRFs were observed in both reactors while 16 *Ex* oTRFs were unique for reactor GR1 (11 unique *Ex* oTRFs of 120, 268, 276, 288, 384, 392, 456, 556, 580, 588 and 624 bp) and GR2 (5 unique *Ex* oTRFs of size 280, 448, 476, 484 and 548 bp). Out of unique *Ex* oTRF in GR1, two (276 and 556 bp) were having RA >1%, while, in GR2 unique *Ex* oTRF 448 and 484 bp were having RA >1% (Fig. A1). In GR1, the *Ex* oTRF 284 bp disappeared at day 84 and reappeared at day 191. *Ex* oTRF 60 and 592 bp disappeared after day 105. The reappearance of *Ex* oTRF 592 bp was observed on day 212 while *Ex* oTRF 60 bp reappeared on day 287. *Ex* oTRF 556 bp was only seen at day105 and 119. The *Ex* oTRF 632 bp was observed (RA >1%) during day 42 and 140-259 (Fig A1A). In control reactor GR2, *Ex* oTRF 60 and 284 bp was observed throughout the operational phase, however, their RA was in the range of 1-13% and 3-15%, respectively. *Ex* oTRF 632 bp was observed in GR2 only in the first three time-points, however, the RA was below 2 % (Fig. A1B). Principal coordinate analysis (PCoA) for the Hpy188III restriction profile showed close clustering of samples (except last three samples) from GR2 while samples from GR1 were scattered under the influence of disturbance which caused an increase in carbon dioxide content (%) and total VFA concentration and eventually resulted in the reduction in methane yield (Fig. A1C).

**A)**

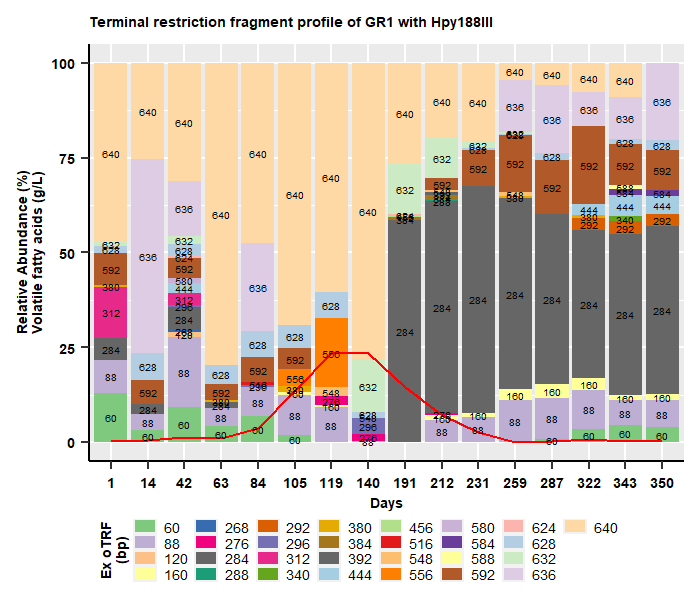

**B)**

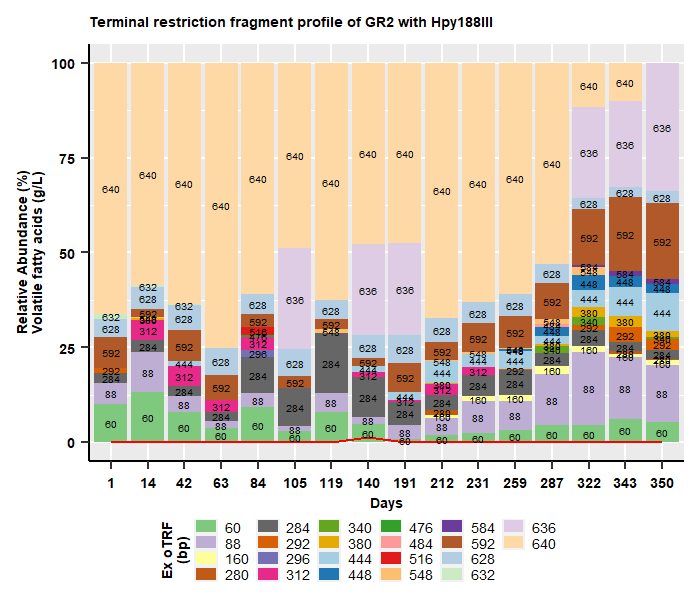

**C)**

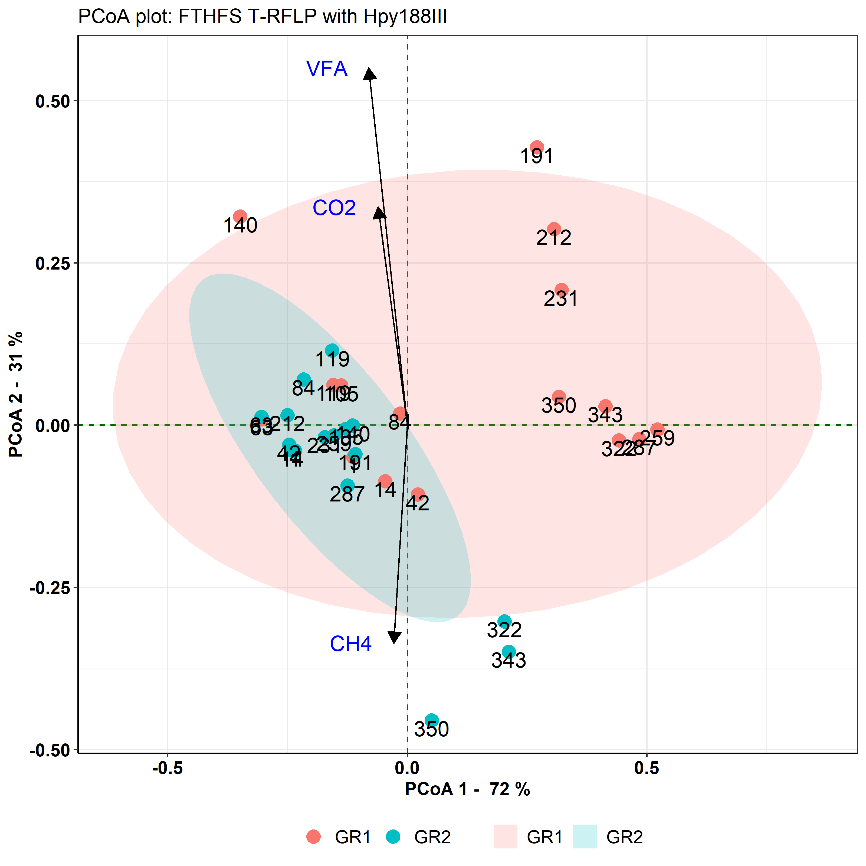

**Figure A1 -** Experimental Terminal restriction fragment length polymorphism (T-RFLP) profile representing *Ex* oTRF with restriction enzyme Hpy188III for **A)** experimental reactor GR1, **B)** control reactor GR2. The red line represents the level of total volatile fatty acids (g/L) at the respective time-points, **C)** Principal Coordinates Analysis (PCoA) plot representing the microbial beta diversity within the experimental (GR1) and control (GR2) reactors using FTHFS T-RFLP profile with Hpy188III. VFA, CH_4_ and CO_2_ are the environmental vectors which represent the level of total volatile fatty acids (g/L), methane content (%) and carbon dioxide content (%), respectively.

#### **16S rRNA gene amplicon sequence data analyses: 16S-community**

16S rRNA gene sequence data analysis showed the presence of 5 phyla with relative abundance (RA) >1% and phylum Bacteroidota, Firmicutes and Cloacimonadota were the most abundant in both GR1 and GR2. Phylum Cloacimonadota disappeared at day 95 and reappeared at day 231 in GR1 (Fig. A2A). At class level 11 classes were observed to have RA >1% and Bacteroidia and Clostridia were the two most abundant classes in both reactors. At the genus level, *Fastidiosipila* (3.6-6.6%) and unknown genus of family Hungateiclostridiaceae (3.5-9.46 %) appeared in GR1 during the whole experimental phase from day 7-125. *Fastidiosiplia* was only seen to have RA >3.5% in GR2 at day 205 (Fig. A2B). An unknown genus of order DTU014 had RA >1% and appeared only from day 231 in GR1, however, it was observed to have a fluctuating presence in GR2 throughout the operational phase. Genus LNR_A2-18 of family Cloacimonadaceae disappeared after day 91 and reappeared at day 231 in GR1 however, it was having fluctuating occurrence in GR2 throughout the operation. Genus *Proteiniphilum* was observed to have increased RA (3-30%) during the same time when total VFA levels were increasing (day 63-140) and with the decrease in total VFA, its RA was also reduced (25-4.6%). No such changes were observed in the control reactor GR2. Unknown genus PeH15 of order Bacteroidales appeared at day 99 with RA ~61% and not seen in next time points from day 105-140, however, it was continuously seen from day 161-259 with RA between 22.4-66.2%. An unknown member of Christensenellaceae R-7 family group was only seen in GR1 at day 161-205 (RA 3.8-6.5%). PeH15 and R-7 group was not seen in GR2 to have RA >3.5%.

Several notable changes happened in GR1 community during the disturbance phase and no similar changes in the community dynamics were observed in the control GR2. This indicates towards the response of the overall community to the increase in the total VFA level, which was also supported by the PCoA and NMDS analysis of GR1 and GR2 (Fig. 3C, Supp. Fig. S7). The unknown genus of family Hungateiclostridiaceae was only seen in GR1 during the disturbance phase and increase in RA of genus *Proteiniphilum* was observed with the increase in VFA. Thus, the appearance and increase in the RA of genus Hungateiclostridiaceae.NA and *Proteiniphilum* can be seen as an indicator of the disturbance. The high RA abundance (~20-60%) of genus PeH15.NA after the disturbance phase suggested towards its involvement in the VFA degradation and during the recovery process. However, according to the current data, we cannot propose if it can be used as an indicator to predict the changes in the community dynamics. Furthermore, a disappearance of phylum Cloacimonadota strongly indicated towards it being an indicator of disturbance. Also, several studies have suggested its role as an indicator in the prediction of disturbance (Calusinska et al., 2018; Klang et al., 2019; Poirier et al., 2020).

**A)**

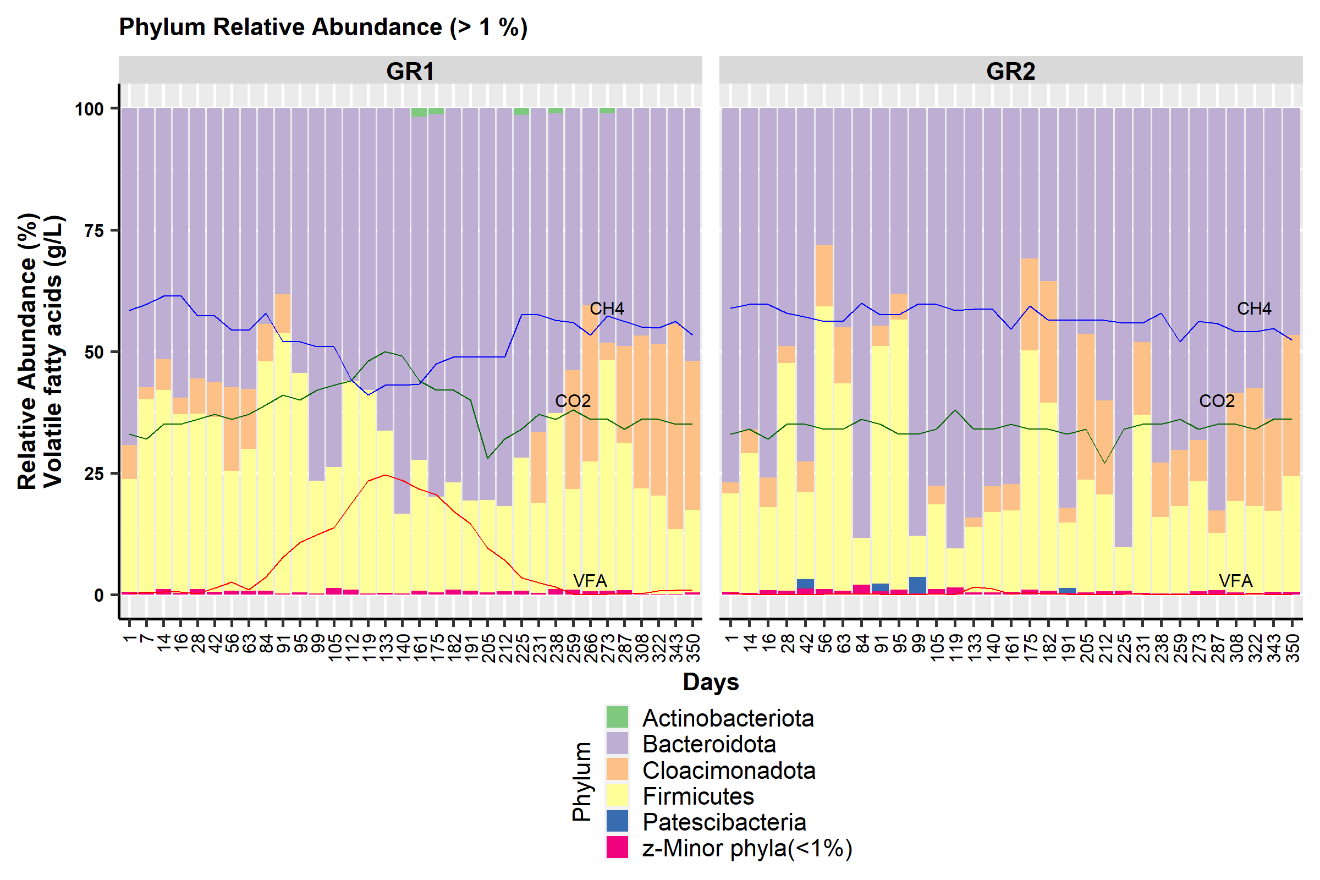

**B)**

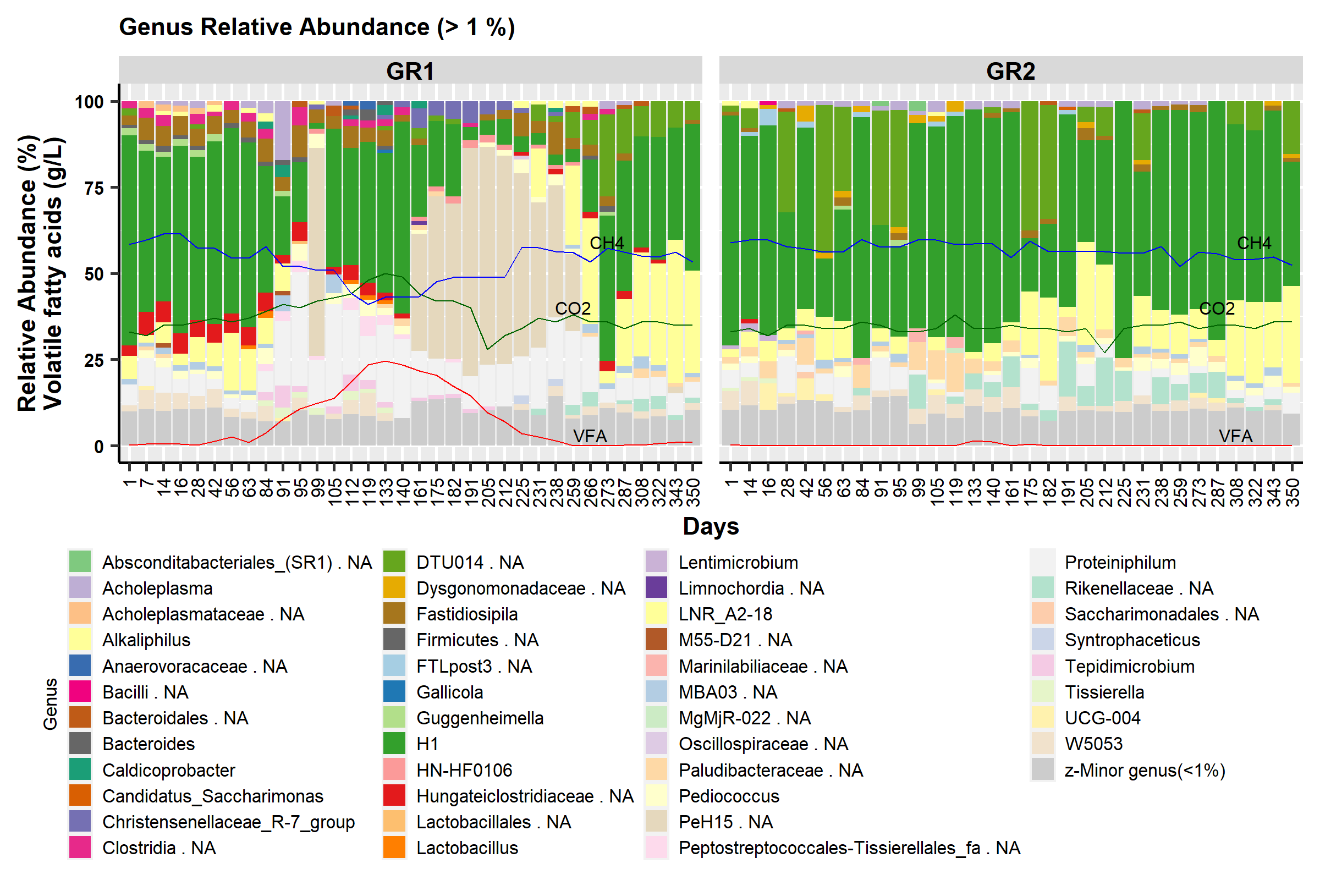

**Figure A2 –** Bar plot representing 16S-community based on 16S rRNA gene amplicons in experimental reactor GR1 and control reactor GR2 at the **A)** phylum level (Relative abundance (RA) >1%) and **B)** genus level (RA >1%). VFA, CH_4_ and CO_2_ represent the level of total volatile fatty acids (g/L), methane content (%) and carbon dioxide content (%), respectively.
