## Supplementary figures for "Profiling acetogenic community dynamics in anaerobic digesters - comparative analyses using next-generation sequencing and T-RFLP"

Running title: Comparative analysis of acetogenic community

Abhijeet Singh^1*^, Bettina Müller^1^, Anna Schnürer^1*^

^1^Anaerobic Microbiology and Biotechnology Group, Department of Molecular Sciences, Swedish University of Agricultural Sciences, Almas Allé 5, Uppsala, SE-750 07, Uppsala, Sweden

**A)**

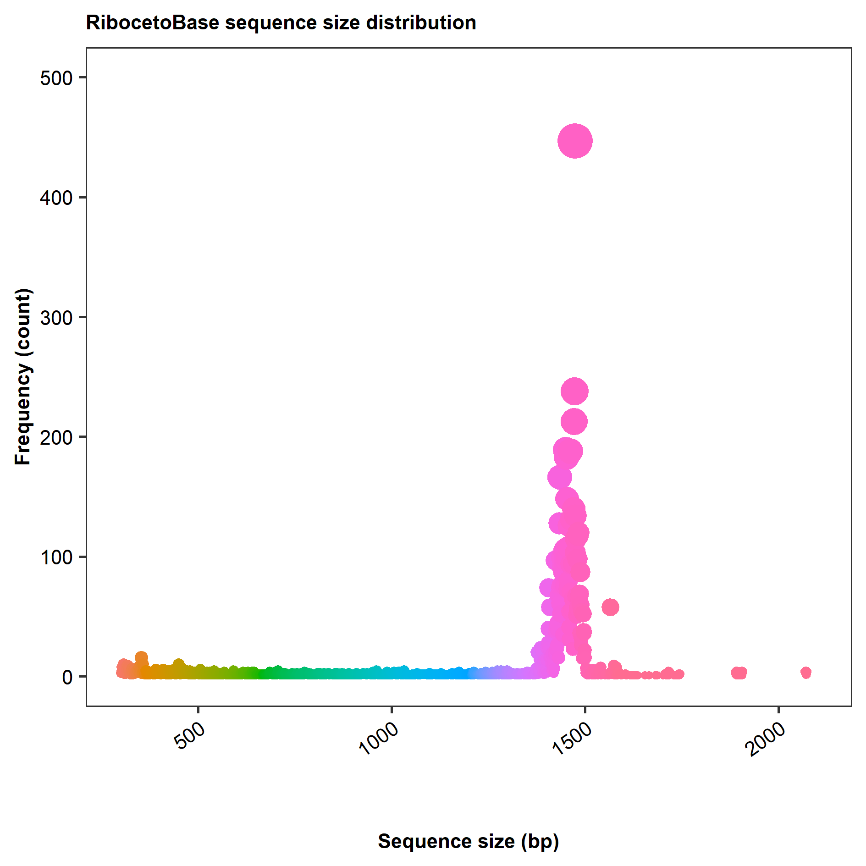

**B)**

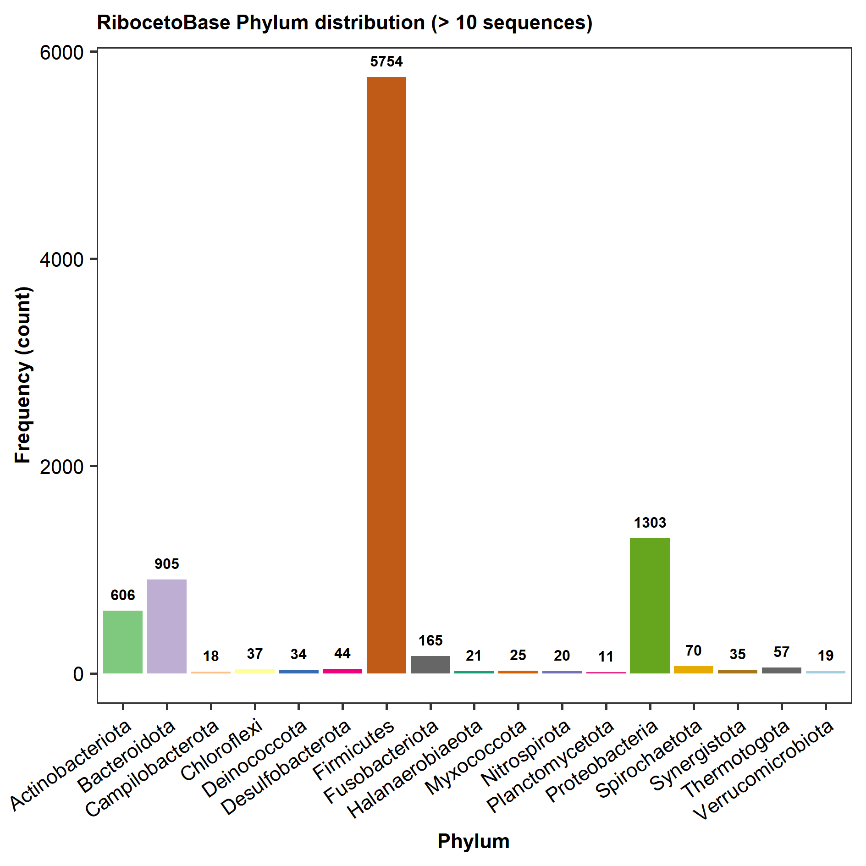

**Supp. Figure S1** - Plots describing the distribution of RibocetoBase sequence according to **A)** lengths **B)** associated taxonomy

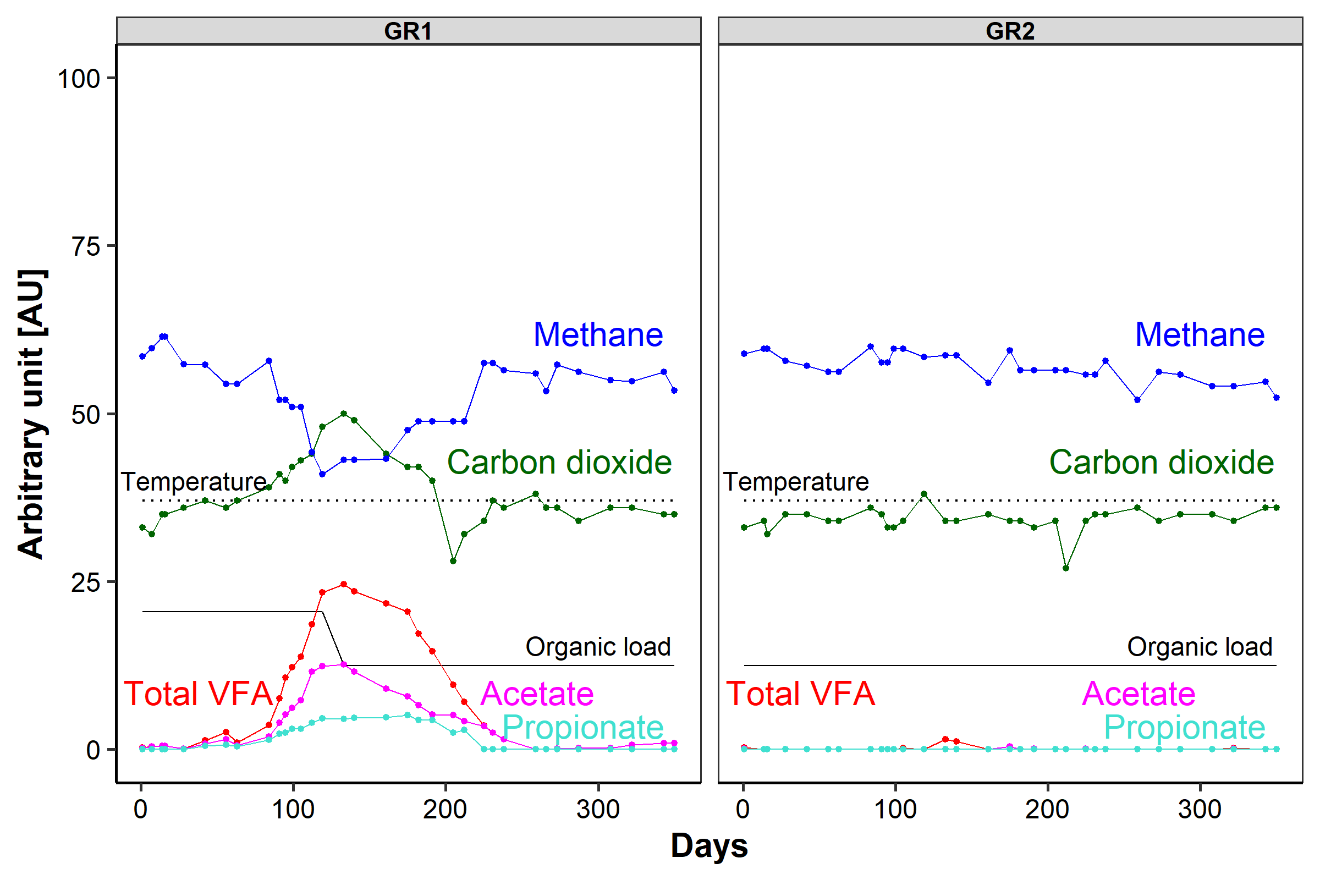

**Supp. Figure S2** - Line chart representing reactor performance of the experimental (GR1) and control (GR2) reactor showing the content of methane (%) and carbon dioxide (%) and concentration of total volatile fatty acid (VFA), acetate and propionate concentration (g/L). The black solid line represents the organic load (20.47 & 12.45 g VS/day) and dotted line represent the operating temperature (37 °C) of the reactors.

**A)	B)**
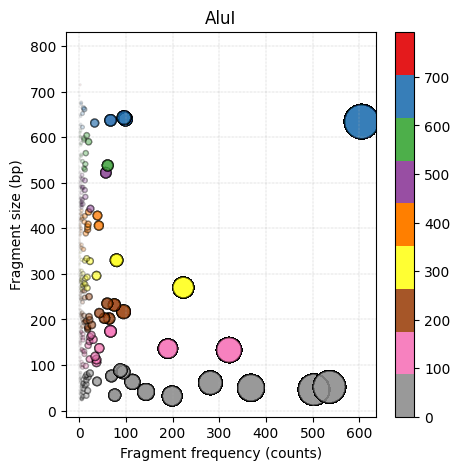

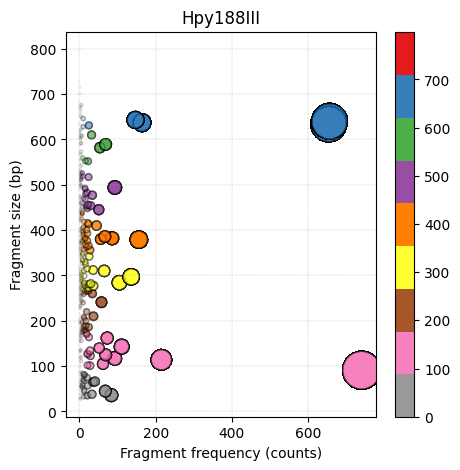

**Supp. Figure S3** - Scatter Plot of *in silico* terminal restriction fragments from AcetoBase FTHFS reference dataset generated by the REDigest program with the restriction enzyme **A)** AluI and **B)** Hpy188III.

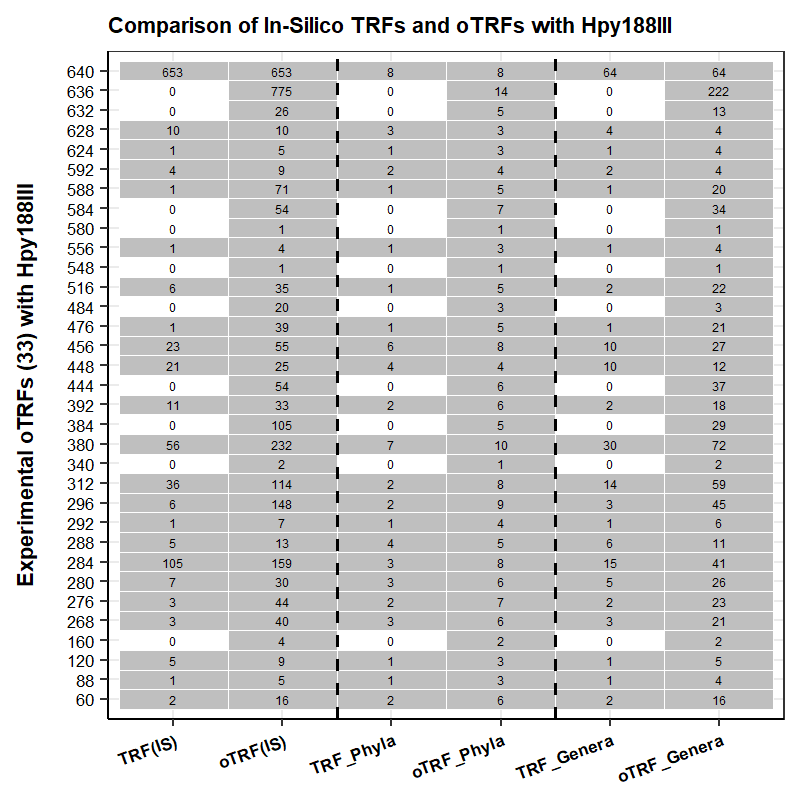

**Supp. Figure S4** - A tabular plot representing FTHFS gene T-RFLP profile during comparison of experimental oTRF versus *in silico* TRF, *in silico* oTRF and count of taxa (phyla and genera) with restriction enzyme Hpy188III.

**A)	B)**

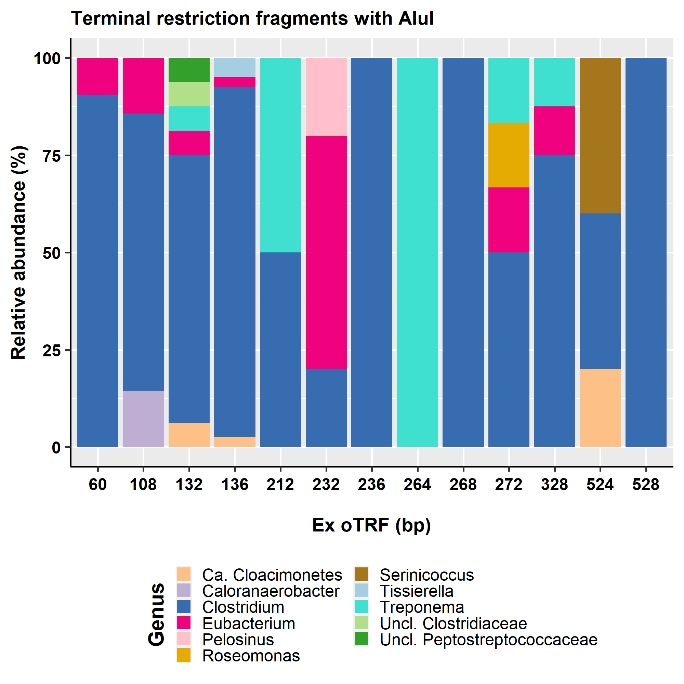

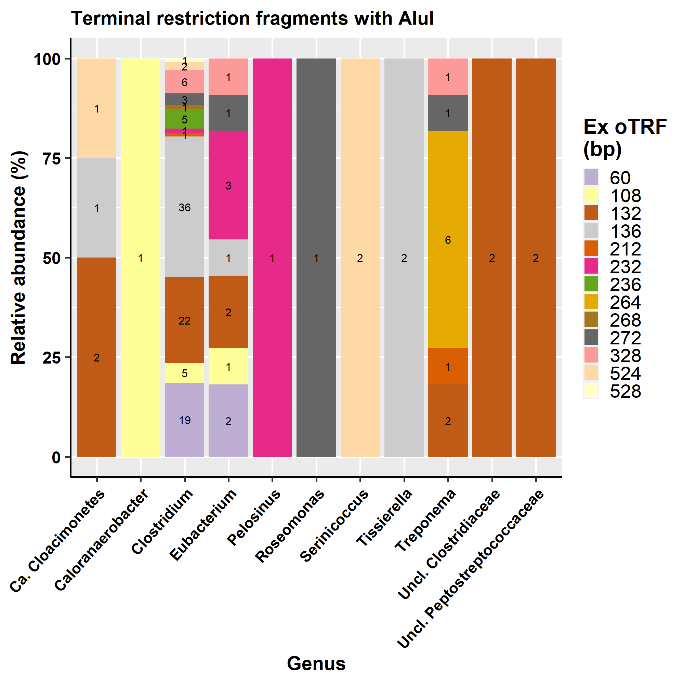

**Supp. Figure S5** - Bar plot representing the taxonomic predictions of the major *Ex* oTRFs generated with the restriction enzyme AluI. A) show the relative abundance of different genus representing the respective *Ex* oTRFs, B) indicate the count based relative abundance of *Ex* oTRFs represented by respective genus in the T-RFLP profile.

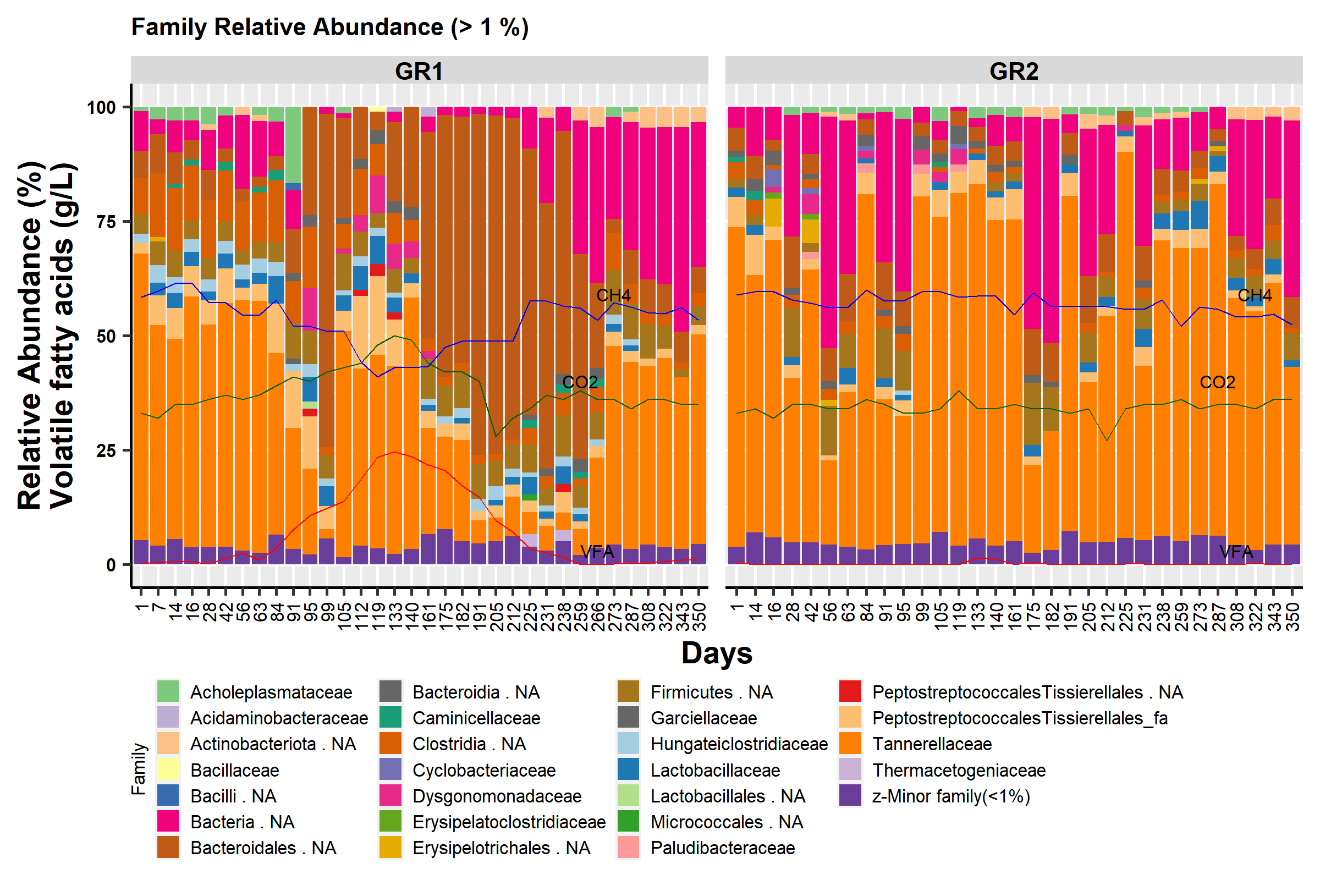

**Supp. Figure S6** - Bar plot representing Riboceto-community in experimental reactor GR1 and control reactor GR2 at the family level (Relative abundance (RA) >1%). VFA, CH_4_ and CO_2_ represent the level of total volatile fatty acids (g/L), methane content (%) and carbon dioxide content (%), respectively.

**A	B**

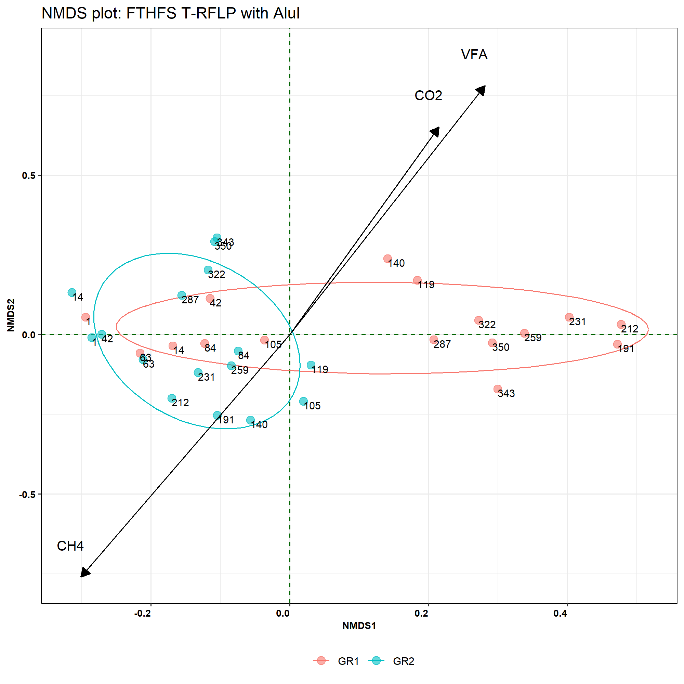

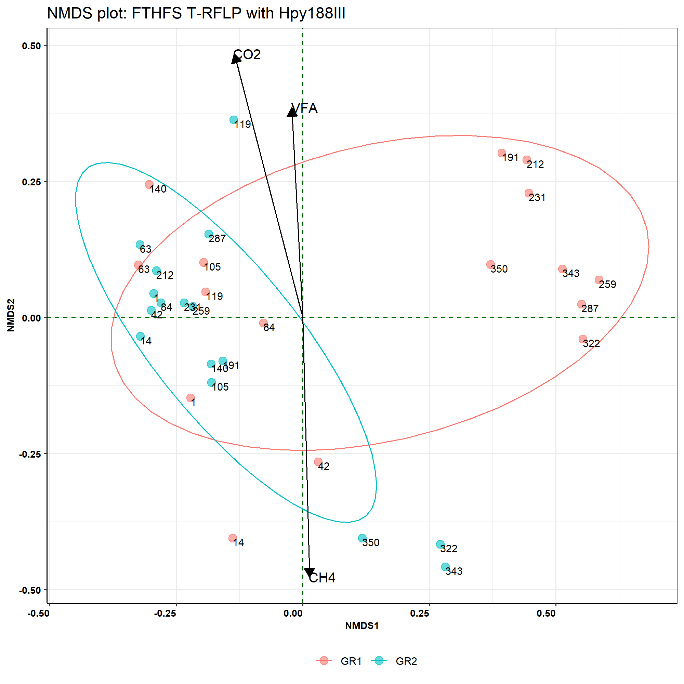

**C	D**

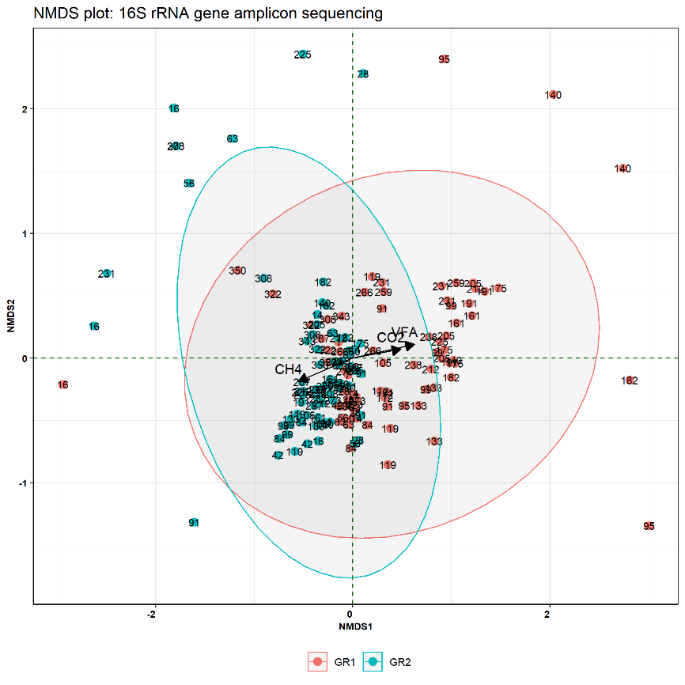

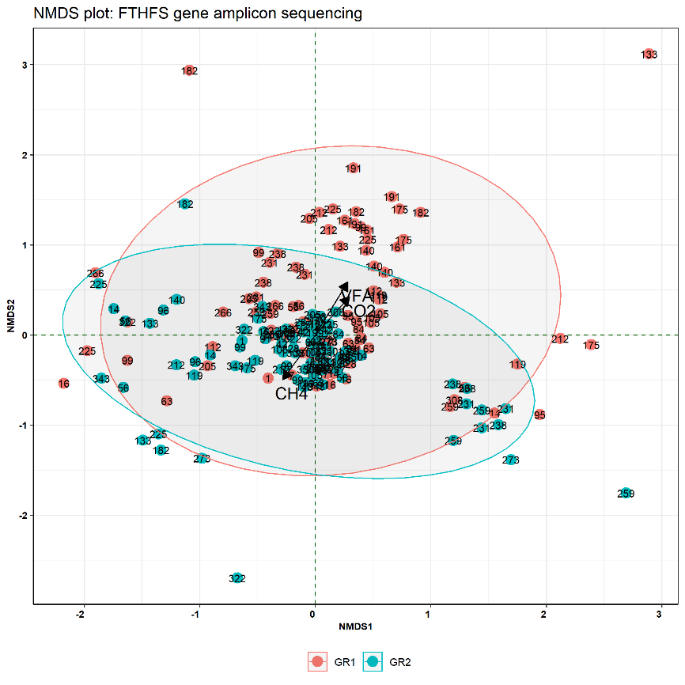

**Supp. Figure S7** - Non-metric Multidimensional Scaling (NMDS) plot representing the microbial beta diversity within the experimental (GR1) and control (GR2) reactors using FTHFS gene T-RFLP profile with restriction enzyme **A)** AluI and **B)** Hpy188III. NMDS plot for the Next-generation sequencing of **C)** 16S rRNA gene amplicons and **D)** FTHFS gene amplicons.

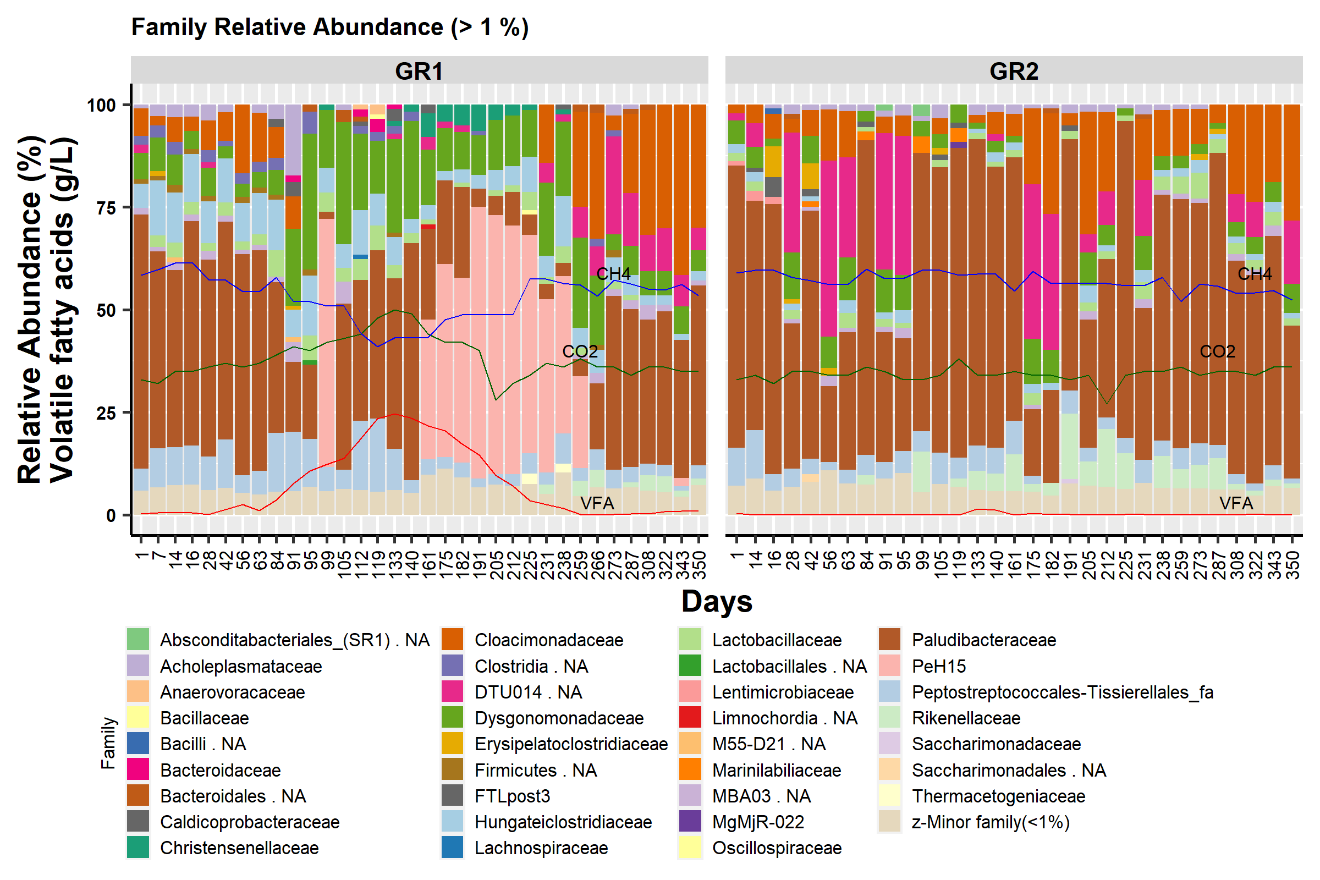

**Supp. Figure S8** - Bar plot representing 16S-community in experimental reactor GR1 and control reactor GR2 at the family level (Relative abundance (RA) >1%). VFA, CH4 and CO2 represent the level of total volatile fatty acids (g/L), methane content (%) and carbon dioxide content (%).

**A)**

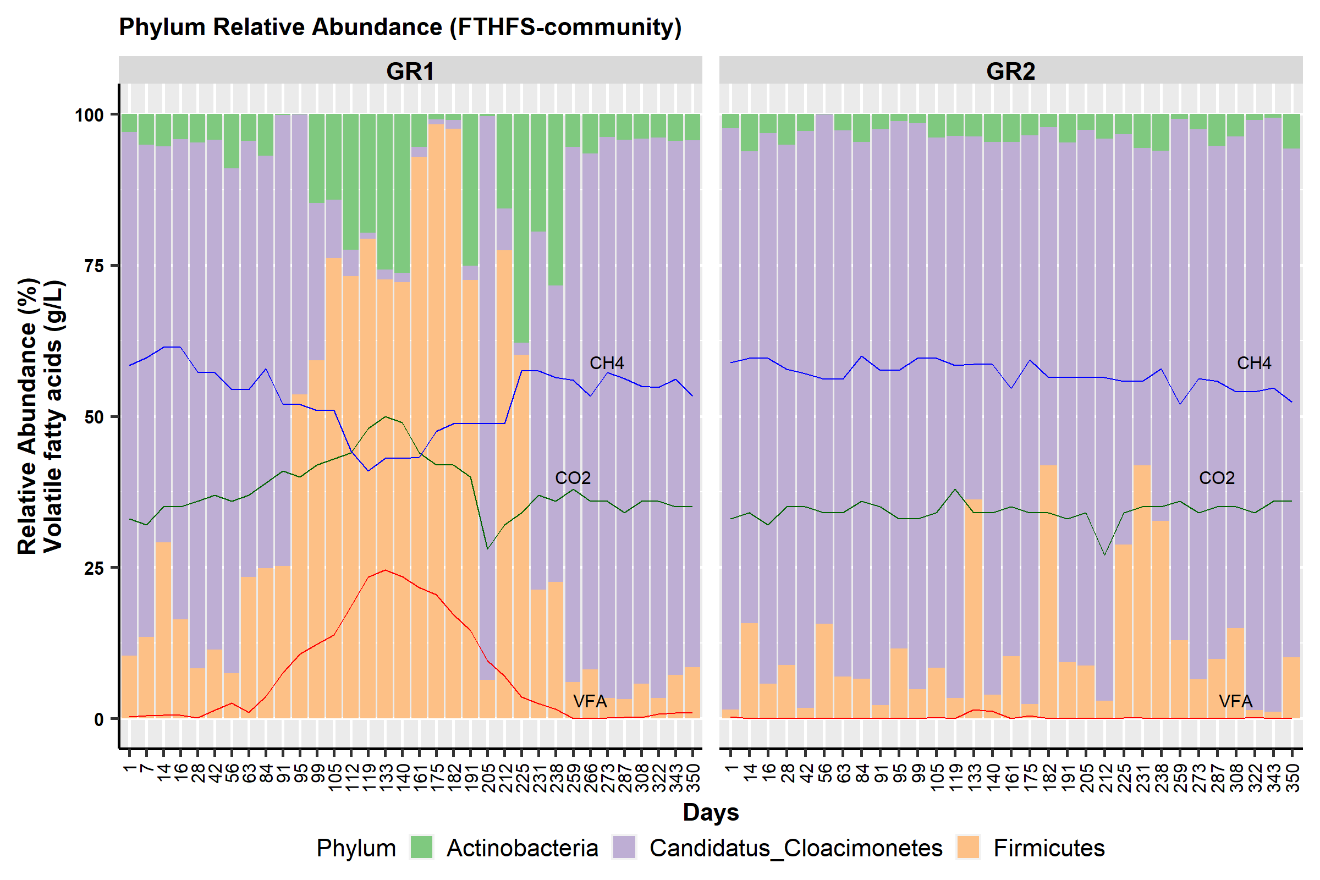

**B)**

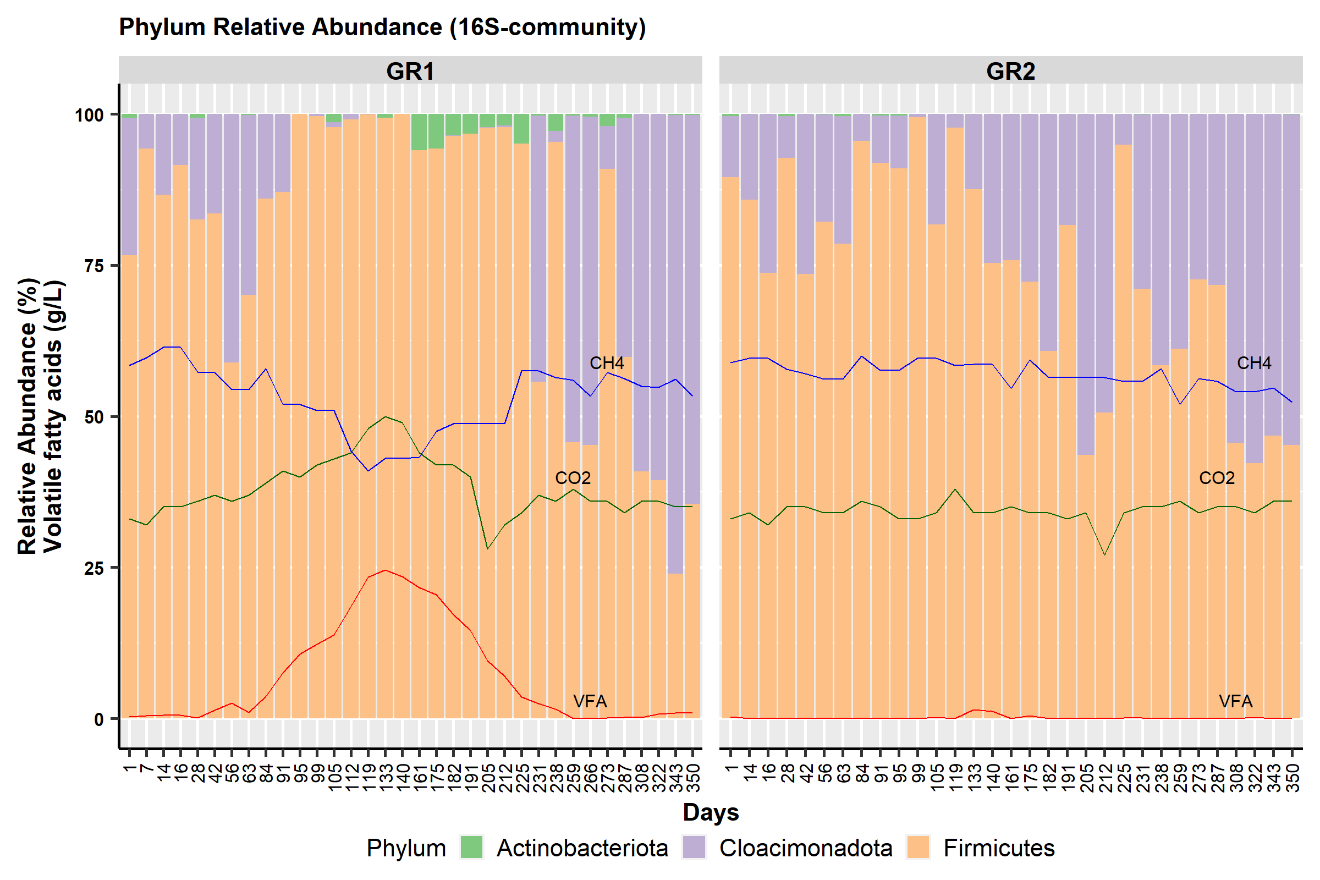

**C)**

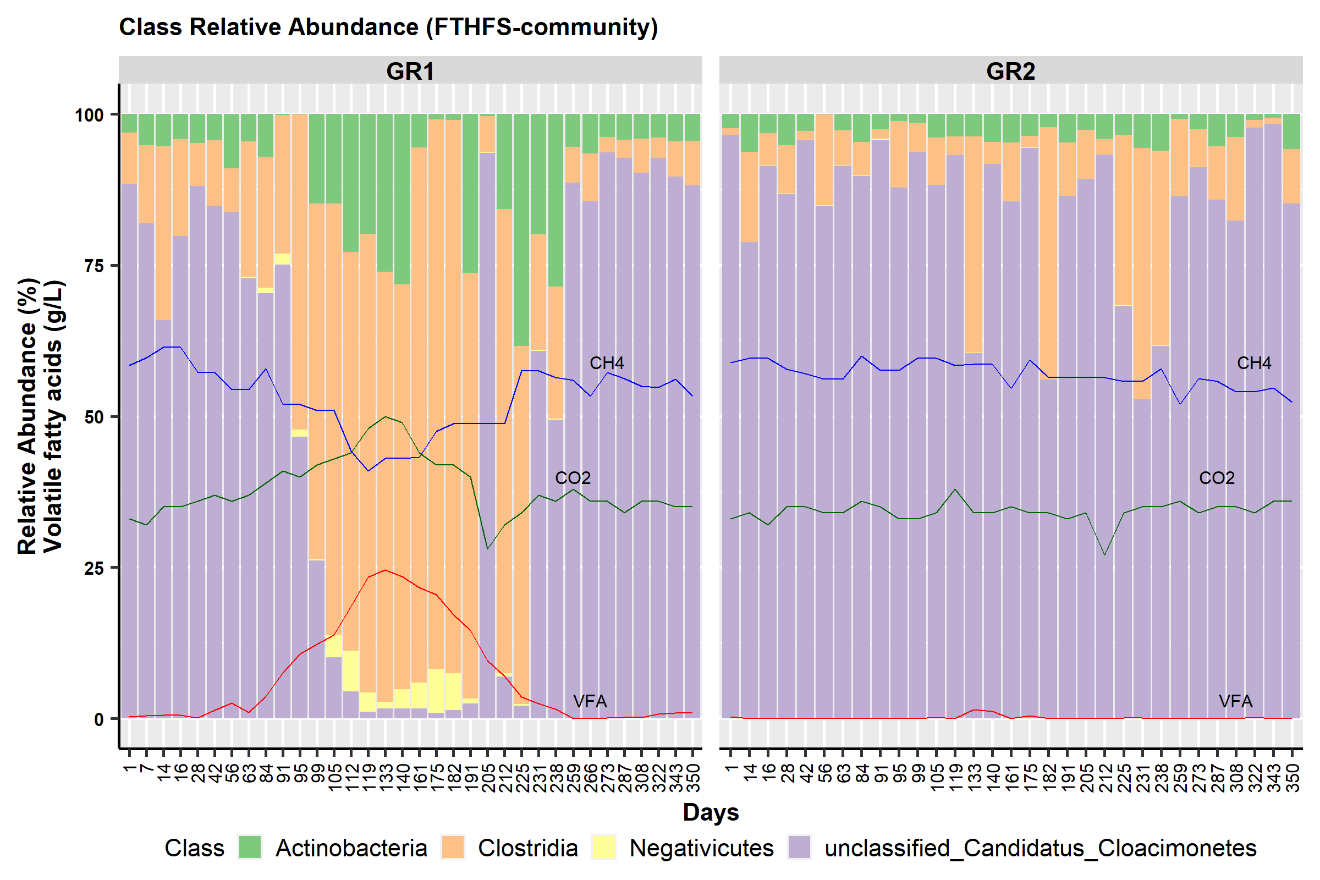

**D)**

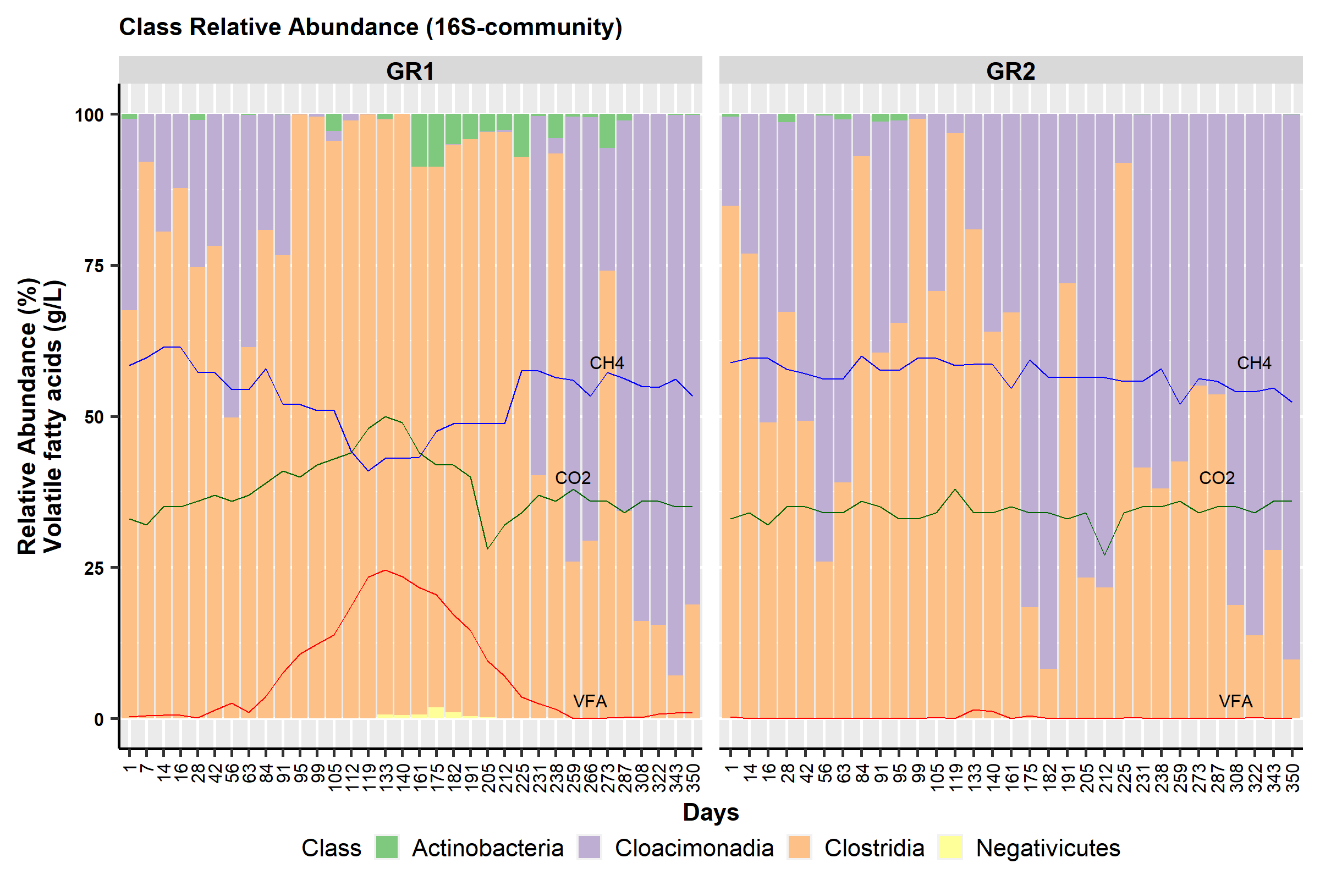

**Supp. Figure S9** - Bar plot representing the top **A)** and **B)** common phyla of FTHFS- and 16S-community, respectively. **C)** and **D)** common classes of FTHFS- and 16S-community, respectively.
